# Structure-based design of soluble prefusion-stabilized herpes simplex virus type 2 glycoprotein B antigens

**DOI:** 10.64898/2026.01.26.701809

**Authors:** Madeline R. Sponholtz, Daphne Ma, Patrick O. Byrne, Yi Shu, Christopher Warren, Nithya Thambi, Ryan S. McCool, Matthew D. Slein, Nicole V. Johnson, William A Rose, Lan Zhang, Eberhard Durr, Dai Wang, Jason. S. McLellan

## Abstract

Herpes simplex virus type 2 (HSV-2) causes genital herpes through latent infection and periodic reactivation. Although antivirals alleviate symptoms, no prophylactic or therapeutic vaccines have been licensed. Glycoprotein B (gB) is a class III fusion protein that mediates entry by irreversibly transitioning from a metastable prefusion conformation to a stable postfusion conformation. Leveraging prior structure-based designs for human cytomegalovirus (HCMV) gB, we engineered amino acid substitutions in HSV-2 gB to stabilize its prefusion conformation. Cryo-EM of the engineered construct revealed a prefusion conformation and non-native dimers of gB trimers. Introduction of N-linked glycosylation sites resulted in the gB-G3 variant, which exhibited improved expression and reduced dimerization. Cryo-EM of gB-G3 bound to neutralizing antibodies yielded a 2.8 Å resolution structure of the stabilized prefusion conformation in a closed state, which differs from the open states observed in recently published HSV gB structures. Both prefusion and postfusion gB variants were evaluated for immunogenicity in mice, delivered either as a protein subunit or mRNA vaccine. Each gB conformation elicited robust humoral and cellular responses. However, prefusion stabilization of gB did not improve neutralizing antibody titers relative to the postfusion construct, consistent with prior observations for HCMV gB. Collectively, these findings reveal insights into prefusion gB conformational dynamics, provide stabilized reagents for studying gB-directed immune responses and inform HSV-2 vaccine design.

## Introduction

Herpes simplex virus type 2 (HSV-2), also known as *Human alphaherpesvirus 2* (*1*), is one of the most prevalent sexually transmitted pathogens, with approximately 13% of adults infected worldwide (*2*). HSV-2 primarily infects mucosal epithelial cells prior to establishing life-long latency in sensory neurons. HSV-2 infection is characterized by alternating latent and lytic cycles that may lead to recurrent genital lesions. Upon reactivation, HSV-2 virions travel from sensory ganglia to reinfect dermal cells at the site of the primary infection, which is typically localized to the genital region. Although many infections are asymptomatic, HSV-2 can cause painful blisters and has been linked to increased risk of HIV-1 acquisition (*3*). HSV-2 can also be transmitted vertically during childbirth, producing neonatal herpes that may progress to life-threatening disseminated disease in infants (*4*). Moreover, chronic infection imposes substantial psychosocial burdens, including stigma and emotional distress (*5*), and emerging studies suggest possible links between HSV-2 and neurodegenerative conditions such as Alzheimer’s disease (*6*). Recurrent outbreaks can be managed with antivirals such as acyclovir (*4*), but there are currently no licensed vaccines approved to treat or prevent HSV-2 infection.

HSV-2 entry requires a coordinated cascade of envelope glycoproteins (*7*). Glycoprotein D (gD) promotes entry by engaging one of three receptor classes: herpesvirus entry mediator (HVEM), nectin-1 or −2, or 3-O-heparan sulfate (*7*). The gH/gL heterodimer is thought to regulate fusion by transmitting the signal from receptor binding to trigger conformational rearrangement of glycoprotein B (gB), a conserved class III fusion protein essential for entry. Upon activation, gB transitions from an initial metastable prefusion conformation to a highly stable postfusion state, a process that drives fusion of the viral and cellular membranes (*7*).

Past HSV protein subunit vaccine efforts have targeted gD or combined glycoprotein antigens. They have yet to fully leverage gB as an antigen, in part due to manufacturing challenges stemming from aggregation and conformational instability, as well as an incomplete understanding of the mechanisms by which gB-specific antibodies neutralize virus infection. When produced recombinantly, gB predominantly adopts the thermodynamically stable postfusion conformation (*8*), raising uncertainty as to whether neutralization-sensitive epitopes present in the native prefusion state are altered or lost. Stabilizing gB in the prefusion conformation may therefore improve functional antibody responses both quantitatively and qualitatively.

Structure-guided stabilization of viral fusion proteins in the prefusion conformation has been transformative for vaccine development targeting class I fusion proteins (*9–12*), leading to the development of vaccines for respiratory syncytial virus (RSV) (*13, 14*). Such stabilization preserves neutralization-sensitive epitopes and elicits higher levels of neutralizing antibody responses (*15*). However, extending this strategy to class III fusion proteins, such as prefusion gB derived from human cytomegalovirus (HCMV) (*16*), Epstein-Barr virus (EBV) (*17*), or HSV-1 (*18*), has not yet resulted in superior neutralization titers compared to the postfusion conformation, although the immunogenicity experiments to date have only assessed protein subunit vaccines and not other delivery modalities, such as mRNA.

Here, we describe a structure-guided design strategy to generate a soluble, prefusion HSV-2 gB ectodomain immunogen. Building on principles established for HCMV gB stabilization, we combine structure-based design with biochemical and biophysical characterization to evaluate prefusion stability and antigenicity. We also assess the immunogenicity of protein subunit and mRNA vaccine formulations of prefusion and postfusion HSV-2 gB. This work advances structure-based vaccine design for class III fusion proteins, potentially paving the way toward an effective HSV-2 vaccine.

## Results

### Design and characterization of single-substitution HSV-2 gB variants

We designed a soluble base construct (gB-B0) comprising the HSV-2 gB ectodomain (HG52 strain, residues 1–731) followed by a C-terminal T4 fibritin (foldon) trimerization motif, an octa-histidine tag, and a Twin-Strep affinity tag. To improve expression, we also removed the proline-rich region following the signal sequence (Δ26–73) (**Figure 1A**). We used this construct, gB-B1, as the base for evaluating all substitutions. To confirm structural homogeneity of our base construct, we performed negative-stain electron microscopy (ns-EM) on gB-B1 in complex with the antigen-binding fragments (Fabs) from murine antibodies 2c, HSV010-13, and BMPC-23 (*19–22*). The resulting 2D class averages indicated that gB-B1 adopts the postfusion conformation, closely resembling previously determined crystal and cryo-EM structures of postfusion HSV-1 gB (*8*) and HSV-2 gB (*23*), respectively (**Supplementary Figure S1**).

**Figure 1:**
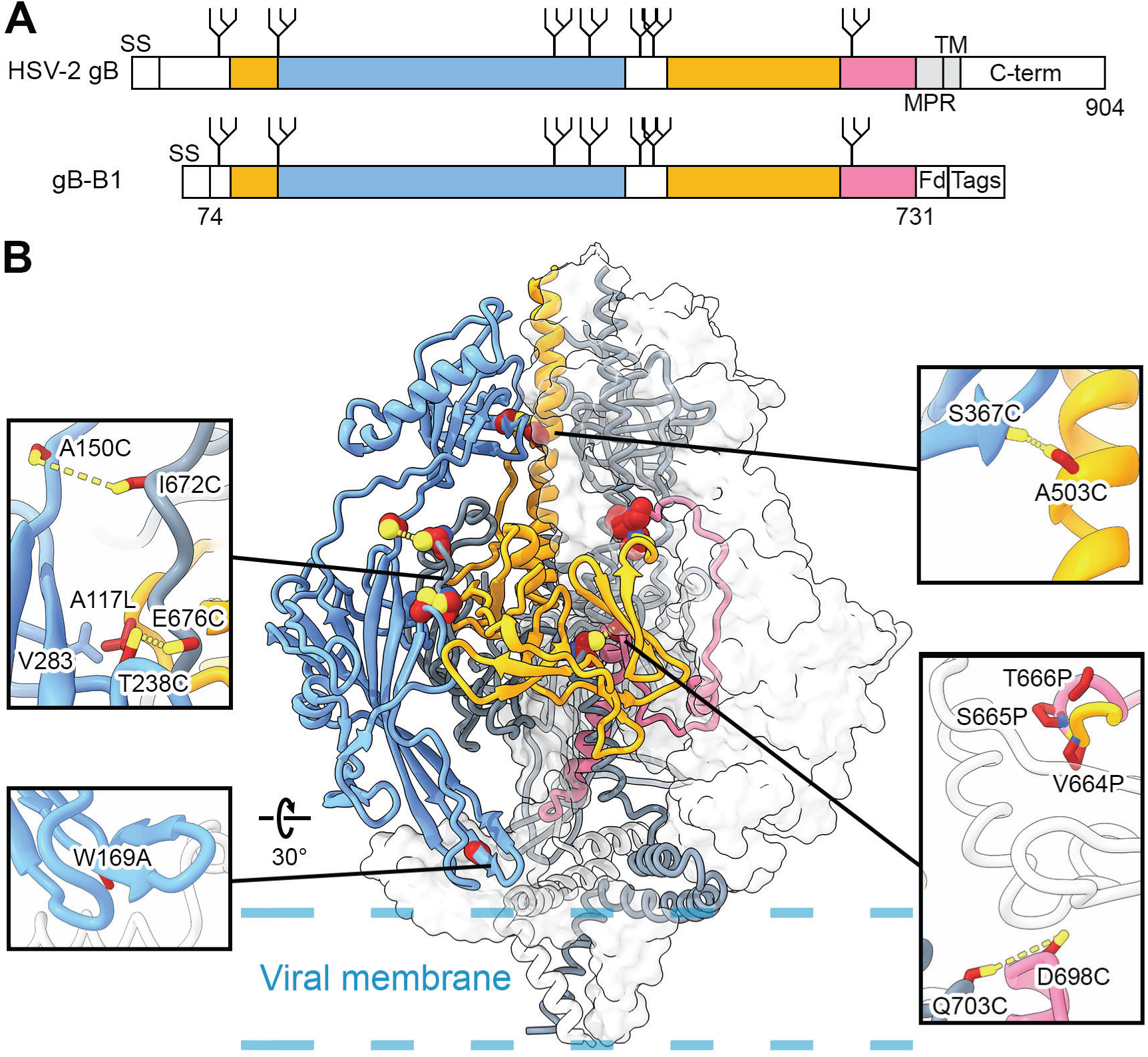
Single substitutions for HSV-2 gB stabilization. (**A**) Schematic of WT HSV-2 strain HG52 gB and ectodomain base construct (gB B1). N-linked glycosylation sites are shown as branched lines. The N-terminal signal sequence (SS) and C-terminal domain (C-term) are shown as white boxes. The first and second regions expected to move during the conformational rearrangement from pre-to-postfusion gB are colored blue and pink, respectively, and the region that does not undergo substantial rearrangement is colored yellow. The MPR and TM domains are shown in gray. The mature gB-B1 construct consists of residues 74-731 followed by the foldon (Fd) trimerization domain and C-terminal tags. (**B**) Side view of trimeric prefusion HSV-2 gB homology model. One protomer is colored as in (A) and shown as a ribbon. One protomer is colored gray and shown as a trace of the α-carbon backbone, and one protomer is shown as a transparent white surface. Single substitutions are shown as spheres with sulfur atoms in yellow, nitrogen atoms in blue, and oxygen atoms in red. The approximate location of the viral membrane is shown as dashed blue lines. Insets show single substitutions as sticks. Anticipated disulfide bonds are shown as yellow dashed lines.

To stabilize the soluble HSV-2 gB ectodomain in the prefusion conformation, we leveraged effective strategies from our recent efforts to stabilize HCMV gB (*16*). We first prepared a homology model of prefusion HSV-2 gB using SWISS-MODEL (*24*) and a rebuilt and re-refined version of the full-length, detergent-solubilized prefusion structure of HCMV gB as the template (*25*) (PDB ID: 7KDP) (**Figure 1B**). From this, we designed six constructs containing substitutions homologous to those previously found to boost expression and stabilize the prefusion conformation of HCMV gB. We also designed five additional HSV-2 gB variants based on the same foundational structure-based design strategies (*26*). In total, we selected 11 designs: 3 proline substitutions, one cavity-filling substitution, six engineered disulfides (five interprotomeric and one intraprotomeric), and one hydrophobicity-reducing substitution to express and purify for characterization. We purified the secreted gB ectodomain variants from 40 mL cultures of FreeStyle 293-F cells to evaluate expression and disulfide bond formation. HSV-2 gB-B1 expression yielded 13.4 mg/L of medium, and the variants yielded between 0.7 and 22.5 mg/L (**Figure 2A**). Only one of the 11 variants tested (W169A) markedly increased expression relative to gB-B1 (∼1.7-fold). Reducing SDS-PAGE analysis of purified variants revealed a predominant band around 110 kDa (**Figure 2B**), consistent with the molecular weight of the glycosylated monomeric HSV-2 gB Δ26–73 ectodomain. On non-reducing SDS-PAGE, three interprotomer disulfide variants (T238C/E676C, A150C/I672C, and D698C/Q703C) yielded bands at ∼460 kDa (**Figure 2B**, black arrow), indicating the presence of disulfide-linked gB trimers. In contrast, the other variants predominantly ran as monomers (**Figure 2B**, white triangle).

**Figure 2:**
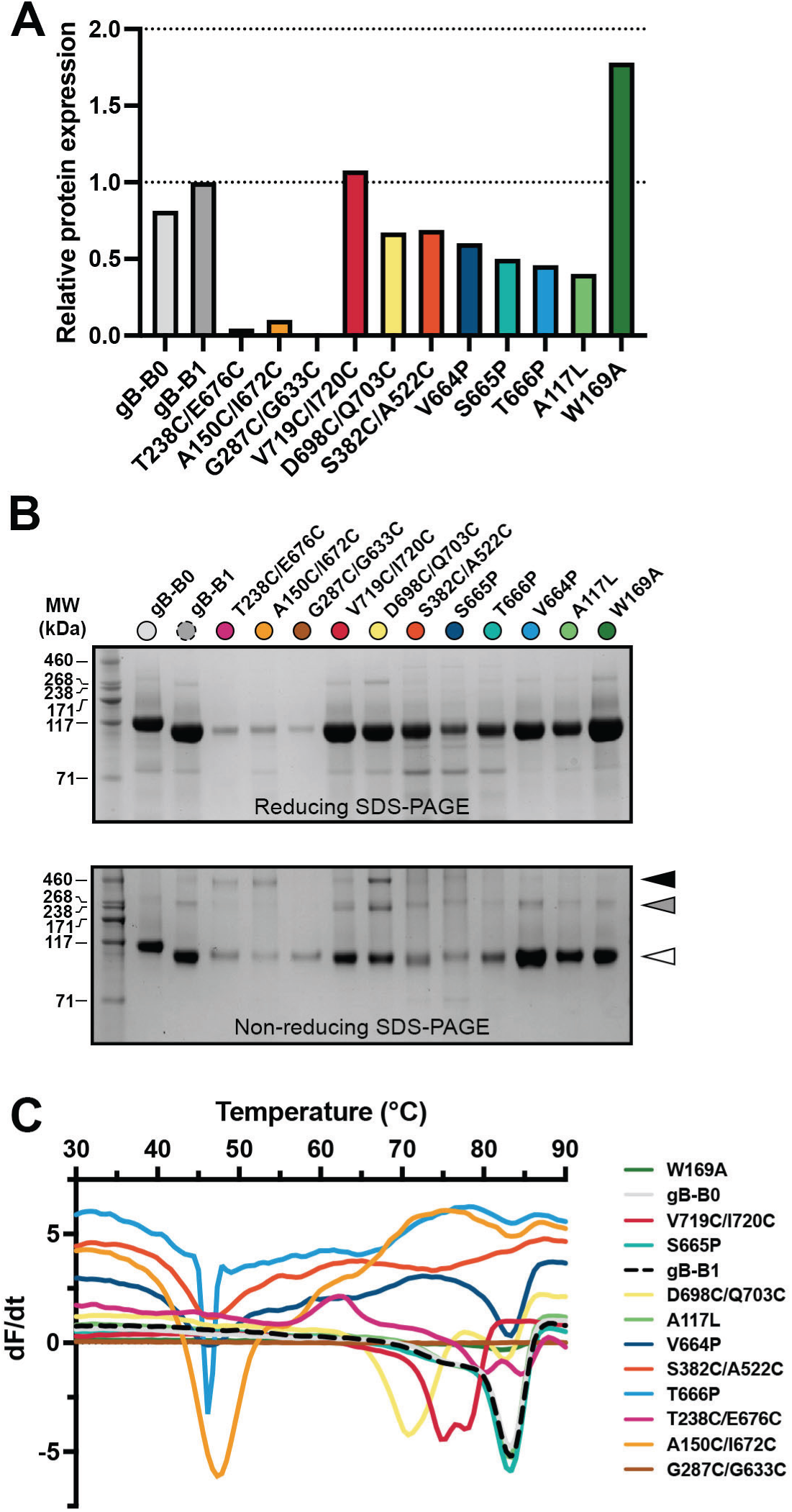
Characterization of single-substitution HSV-2 gB variants. (**A**) Relative expression levels of individual variants. (**B**) Reducing and non-reducing SDS-PAGE of gB variants. Molecular weight standards are indicated at the left in kDa. White, gray, and black triangles correspond to monomeric, dimeric, and trimeric gB molecular weights, respectively. (**C**) Differential scanning fluorimetry (DSF) analysis of gB variant thermostability.

Variants were then analyzed by differential scanning fluorimetry (DSF) to evaluate thermal stability. When evaluated by DSF, gB-B1 exhibited a Tm of 83 °C (**Figure 2C**). Although a subset of variants had DSF profiles that closely resembled those of gB-B1, several displayed distinct characteristics. Some variants displayed two thermal transitions: one at a lower temperature (∼45 °C) and one at 83 °C. Given the high stability of the postfusion conformation, we hypothesized that the ∼45 °C transition corresponds to prefusion-stabilized trimers. Excluding G287C/G633C, which did not yield a clear melting temperature, all disulfide variants (T238C/E676C, A150C/I672C, V719C/I720C, D698C/Q703C, and S382C/A522C) as well as the proline variants V664P and T666P exhibited reduced melting temperatures relative to gB-B1. Overall, no variants exhibited uniformly improved characteristics relative to gB-B1 across all biochemical assays, but several disulfide (T238C/E676C, A150C/I672C, V719C/I720C, and D698C/Q703C) and proline (V664P, S665P, and T666P) variants exhibited some favorable characteristics (**Figure 2**).

### Biochemical and structural characterization of combination variants

We engineered six HSV-2 gB combination variants (gB-S1–S6; **Table 1**), each incorporating sets of two to four individually beneficial amino acid substitutions, and evaluated them for additive effects on expression and stability. We first selected the interprotomer disulfides T238C/E676C and A150C/I672C—analogous (based on sequence alignment) to the disulfides V134C/I653C and H222C/E657C—which stabilize HCMV gB in a prefusion-like conformation (*16*) and appeared to partially form in HSV-2 gB (**Figure 2B**). These disulfides were designed to bridge DI and DV, preventing them from separating during the transition from the pre- to postfusion conformation. We additionally included two hinge-stabilizing proline substitutions, S665P and T666P, located in DV, which must elongate to pack between the central helices of DIII during the structural conversion (*23*). Although these prolines did not boost gB expression, T666P had a favorable DSF profile and we hypothesized that stabilizing the DV hinge could have additive beneficial effects on prefusion stability when paired with successfully formed disulfides. Of the six combination variants evaluated, all except gB-S6 (T238C/E676C, S665P, T666P) had reduced expression relative to gB-B1 and their individual substitutions (**Figure 3A**). Notably, gB-S6 expressed similarly to gB-B1, indicating that the combination of these substitutions recovered the reduced expression observed for the variants containing each of the individual substitutions. Non-reducing SDS-PAGE analysis also showed that gB-S6 exhibited a substantial increase in trimer species relative to gB-B1, as did gB-S5 (**Figure 3B**, black arrow). DSF analysis revealed that most combination variants exhibited markedly lower melting temperatures than gB-B1 (**Figure 3C**), as expected for prefusion-stabilized constructs. Both gB-S5 and gB-S6 exhibited a Tm of 64 °C, whereas gB-S2 and gB-S3 exhibited slightly lower Tm values of 60 and 62 °C, respectively.

**Figure 3:**
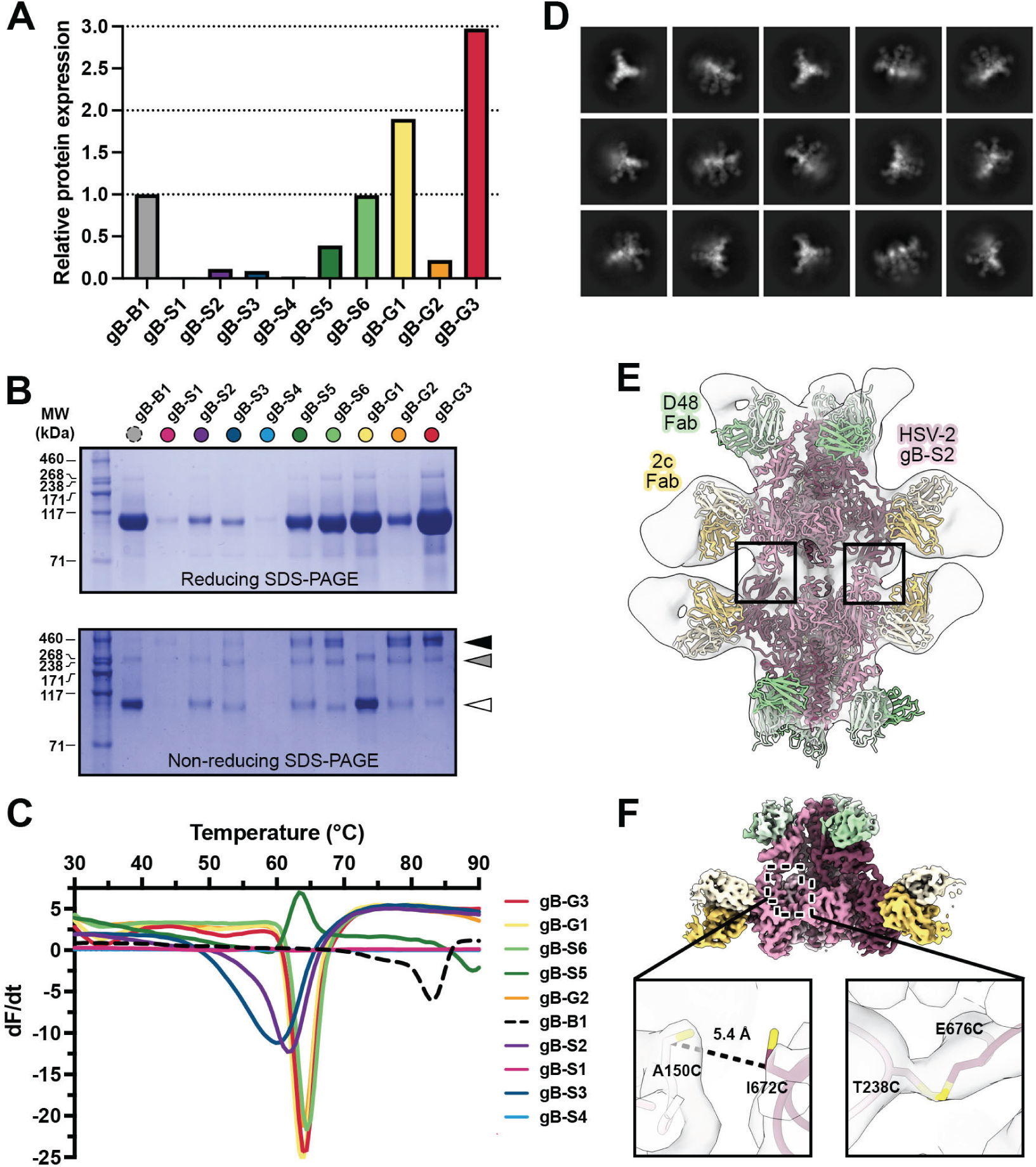
Characterization of combination gB variants. (**A**) Relative expression levels of combination variants. Dashed lines denote 1-fold and 2-fold increases in expression relative to gB-B1. (**B**) Reducing and non-reducing SDS-PAGE gels of gB variants. Molecular weight standards are indicated at the left in kDa. White, gray, and black triangles correspond to monomeric, dimeric, and trimeric gB molecular weights, respectively. (**C**) DSF analysis of gB variant thermostability. (**D**) 2D class averages of gB-S2 in complex with Fabs 2c and D48. (**E**) Medium-resolution 3D reconstruction of non-native dimers formed by the gB-S2-Fab complex. Two copies of a model of the gB-S2-2c-D48 complex are shown as ribbon diagrams and fit into the partially transparent map volume. Black boxes indicate the dimer interface near the gB fusion loops. (**F**) A 3.8 Å resolution reconstruction of the gB-S2-Fab complex, shown with colored map density as in (E). Insets show zoomed-in views of the disulfide substitutions in gB-S2 fit into the partially transparent map volume. Sulfur atoms are colored yellow.

**Table 1.** HSV-2 gB combination variants.

| **Combo**<br>**variant** | **Substitution 1** | **Substitution 2** | **Substitution 3** | **Substitution 4** |
| --- | --- | --- | --- | --- |
| gB-S1 | T238C/E676C | A150C/I672C | - | - |
| gB-S2 | T238C/E676C | A150C/I672C | S665P | T666P |
| gB-S3 | T238C/E676C | A150C/I672GCG | S665P | T666P |
| gB-S4 | T238C/E676C | A150C/N671Q/I672GCG | S665P | T666P |
| gB-S5 | T238C/E676C | S665P | - | - |
| gB-S6 | T238C/E676C | S665P | T666P | - |
| gB-G1 | T238C/E676C | S665P | T666P | M213T |
| gB-G2 | T238C/E676C | S665P | T666P | A256N/F257L/H258T |
| gB-G3 | T238C/E676C | S665P | T666P | H172N/Y174T |

We next conducted cryo-EM studies on a subset of gB combination variants to assess their conformations. To aid the characterization, we formed complexes with two moderate-affinity, neutralizing antibodies, 2c and D48, which bind DI and DII, respectively (*19–21*). Micrographs collected for gB-S2—a combination variant containing two proline substitutions and two disulfide bond substitutions homologous to those in HCMV gB-C7—were of sufficient quality to obtain 2D class averages and an initial moderate-resolution (∼11 Å) 3D reconstruction (**Table 1**, **Figure 3D,E**). We found that gB-S2 is stabilized in the prefusion conformation but forms non-native dimers-of-trimers with an extensive DI interface mediated in part by the hydrophobic residues in the fusion loops (**Figure 3E**). Further processing and refinement with C3 symmetry resulted in a 3.8 Å resolution reconstruction of a single prefusion trimer that enabled analysis of the stabilizing substitutions (**Figure 3F**). Although clear, connected map features were observed for the T238C/E676C disulfide, the map for the A150C/I672C disulfide was disconnected, and the Cβ atoms for these residues were separated by ∼ 5 Å, indicating this disulfide may not have successfully formed or was broken as a result of electron-beam-induced damage (**Figure 3F**). These results suggested that T238C/E676C in combination with S665P and T666P (the substitutions present in gB-S6) may suffice for prefusion stabilization of HSV-2 gB. This observation, combined with the favorable expression and thermal-transition profiles observed for gB-S6, led us to select this variant for additional studies.

To reduce non-native dimerization of gB trimers, we further modified the gB-S6 construct by introducing N-linked glycosylation sites into DI, targeting the dimerization interface observed in the gB-S2 reconstruction (**Figure 3E**). This strategy was previously effective in minimizing oligomerization of postfusion HCMV gB (*27*). We generated three glycosylation variants—gB-G1, gB-G2, and gB-G3—by incorporating N-linked glycosylation motifs at residues 211, 257 (proximal to fusion loop II), and 172 (proximal to fusion loop I), respectively (**Table 1**). Upon purification from 40 mL cultures of FreeStyle 293-F cells, gB-G1 and gB-G3 exhibited increased expression relative to gB-S6, whereas gB-G2 expression was substantially reduced (**Figure 3A**). Non-reducing SDS-PAGE analysis showed a decreased proportion of the trimer species for gB-G1, while gB-G2 and gB-G3 maintained high levels of trimer formation relative to gB-S6 (**Figure 3B**, black arrow). The glycosylation designs appeared to have little-to-no impact on thermostability, as gB-G1, gB-G2, and gB-G3 all exhibited Tm values comparable to gB-S6 at ∼64 °C (**Figure 3C**). Among all variants, gB-G3 demonstrated the most favorable profile, with a three-fold increase in expression compared to gB-B1 (**Figure 3A**), the highest trimer content on non-reducing SDS-PAGE (**Figure 3B**), and a Tm value of 64 °C (**Figure 3C**), supporting its selection for further structural analysis.

### HSV-2 gB-G3 is stabilized in the prefusion conformation

We determined a cryo-EM structure of HSV-2 gB-G3 in complex with Fabs 2c and D48 at a global resolution of 2.8 Å with C3 symmetry refinement (**Figure 4A, Supplementary Table 1**). The final model spans residues 90–721, with residues 168–179, 247–260, and 461–498 omitted due to poor map features for the apex and fusion loops that precluded unambiguous model building. The stabilized HSV-2 gB-G3 structure contains an N-terminal helix (residues 92–100) that packs against DII (**Figure 4B**), a feature observed in one of the recent HSV-1 prefusion-stabilized gB constructs (*28*) but not the other (*18*). We were also able to model residues at the HSV-2 gB membrane-distal apex that adopt extended α-helices, in agreement with our previously determined structure of HCMV gB stabilized in a prefusion-like conformation (*16*). Importantly, the high-resolution map confirmed all sidechains at the sites of substitution and interprotomer disulfide bond formation, as evinced by the contiguous map between the cysteines of the T238C/E676C disulfide (**Figure 4B, Supplementary Figure S2**), and the introduced prolines at positions 665 and 666 both adopted the *cis* conformation.

**Figure 4:**
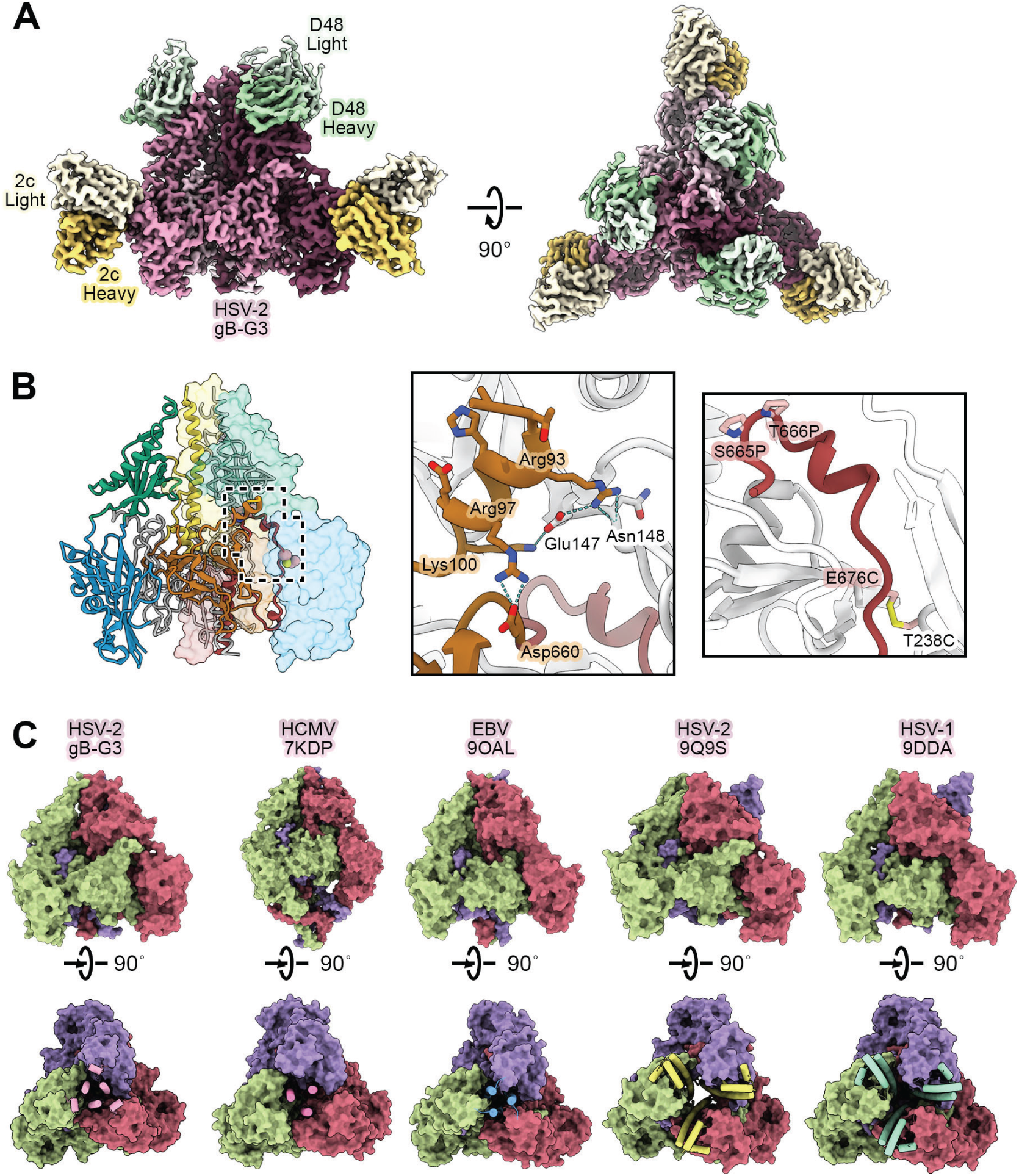
Cryo-EM structure of gB-G3 bound to 2c and D48 Fabs. (**A**) Side (*left*) and top (*right*) views of the EM map of HSV-2 gB-G3 complexed with 2c and D48 Fabs. 2c heavy chain (HC) is yellow, 2c light chain (LC) is pale yellow, D48 HC is green, and D48 LV is pale green, and gB-G3 is three shades of pink. (**B**) Side view of the gB-G3 complex model (*left*). One protomer of the model is colored by domain DI:Blue, DII: green, DIII: yellow, DIV: orange, DV: red and shown as a ribbon diagram. The second is colored gray and shown as a cartoon tube trace of the α-carbon backbone, and the third is shown as a transparent surface also colored by domain. The first inset (*center*) shows a zoomed view of the N-terminal helix. The second inset (*right*) shows a zoomed view of the substitutions that comprise stabilizing substitutions in gB-G3. In both insets, key residues are shown as sticks, and salt bridges and hydrogen bonds are shown as green dashed lines with disulfide bonds in yellow. (**C**) Side (top row) and top-down (bottom row) views of surface renderings of several prefusion-stabilized herpesvirus gB trimers with PDB codes listed below each label. The central helices are shown as cylinders in the top-down views, with their molecular surfaces omitted for clarity.

The overall architecture of HSV-2 gB-G3 closely resembles that of recently reported prefusion-stabilized HSV gB structures (*18, 28*), as well as prefusion gB structures from HCMV (*25*) and EBV (*17*) (**Figure 4C**). Among prefusion gB structures, the principal differences lie in the positioning of the central helices and the degree to which the membrane-distal trimer apex adopts an open or closed conformation. The central helices of HSV-2 gB-G3 converge at the apex, yielding a closed conformation that closely resembles both the cross-linked full-length wildtype HCMV gB structure (*25*) and a recently determined prefusion-stabilized EBV gB ectodomain structure (*17*). In contrast, the prefusion-stabilized HSV-1 gB structures (*18, 28*) exhibit splayed central helices at the apex, resulting in an open conformation. This open conformation was also observed in a structure of full-length, wild-type HSV-2 gB in complex with a prefusion-specific nanobody that was elicited by immunization with the stabilized HSV-1 gB protein (*28*). Interestingly, in a preprint describing the stabilization of several herpesvirus gB ectodomains, a prefusion-stabilized HSV-1 gB construct adopted both open and closed forms in the same dataset (*29*). These observations support the notion that the central helices and trimer apex are conformationally dynamic and capable of “breathing,” potentially modulated by stabilizing substitutions, antibody binding, or other factors.

To better understand the unique epitopes that may be presented on the prefusion conformation, we performed a detailed structural comparison of HSV-2 gB-G3 and the postfusion structure of HSV-2 gB (PDB ID: 8RH1) (*23*) (**Supplementary Figure S3**). First, relative solvent-accessible surface area (rSASA) values were calculated for each residue in both conformations, and the difference in rSASA values was used to visualize surface patches that show differential accessibility between the two conformations (**Supplementary Figure S3B**). This analysis demonstrated only a few patches of adjacent residues, primarily located within DI near the fusion loops, that appear significantly more accessible in the prefusion conformation. To better understand changes to the local environment that may not be obvious in the rSASA analysis, we also defined a local ‘neighborhood’ as all residues within 10 Å of each residue in each conformation and compared the similarity of each residue neighborhood using the Jaccard similarity metric (**Supplementary Figure S3C**). A plot of the per-residue SASA difference and neighborhood Jaccard similarity scores further demonstrates that many of the prefusion-specific epitopes primarily reside in DI near the fusion loops (**Supplementary Figure S3D**). Importantly, these areas are located proximal to the viral membrane and would likely not be accessible in native HSV-2 virions. These analyses support findings from other class III viral fusion proteins (*18*) and indicate that prefusion-to-postfusion conformational changes largely occur via multiple rigid-body movements, with few epitopes being exposed in one conformation and buried in the other.

### Immunogenicity of prefusion and postfusion HSV-2 gB proteins in mice

To compare the immunogenicity of prefusion vs postfusion HSV-2 gB proteins *in vivo*, we immunized mice with two doses (0.5 µg or 2 µg) of the HSV-2 gB-B1, gB-S6, and gB-G3 protein ectodomain constructs adjuvanted with alum or a TLR4 agonist and liposome (TLR4+liposome) and examined antibody and T cell responses (**Supplementary Figure S4**). The differences between vaccine antigen groups were more pronounced at day 27 (∼4 weeks post-dose 1) compared to day 42 (2 weeks post-dose 2) (**Figure 5A; Supplementary Table 2**), as all immunization groups, regardless of adjuvant, induced a robust boost in ELISA geometric mean titers (GMTs) after the second dose. Moreover, statistically significant differences were observed only in vaccinations utilizing TLR4+liposome, but not alum, as an adjuvant (**Figure 5A**). Notably, immunization with gB-B1 postfusion constructs induced higher GMTs at day 27 compared to gB-S6 (2.3 to 7.3-fold,) or gB-G3 (12.7 to 51.3-fold) vaccine groups (**Supplementary Table 2**), but statistical significance between GMTs was only reached with TLR4+liposome as an adjuvant (**Figure 5A**). gB-S6 also showed an increase in ELISA GMTs compared to gB-G3 at day 27 with alum adjuvant (2.2-fold for both doses) and TLR4+liposome adjuvant (12.5-fold for the 2 µg dose or 14.7-fold for the 0.5 µg dose), but this effect was not statistically significant and less pronounced than comparisons between gB-B1 and gB-G3 immunization groups (**Supplementary Table 2**).

**Figure 5.**
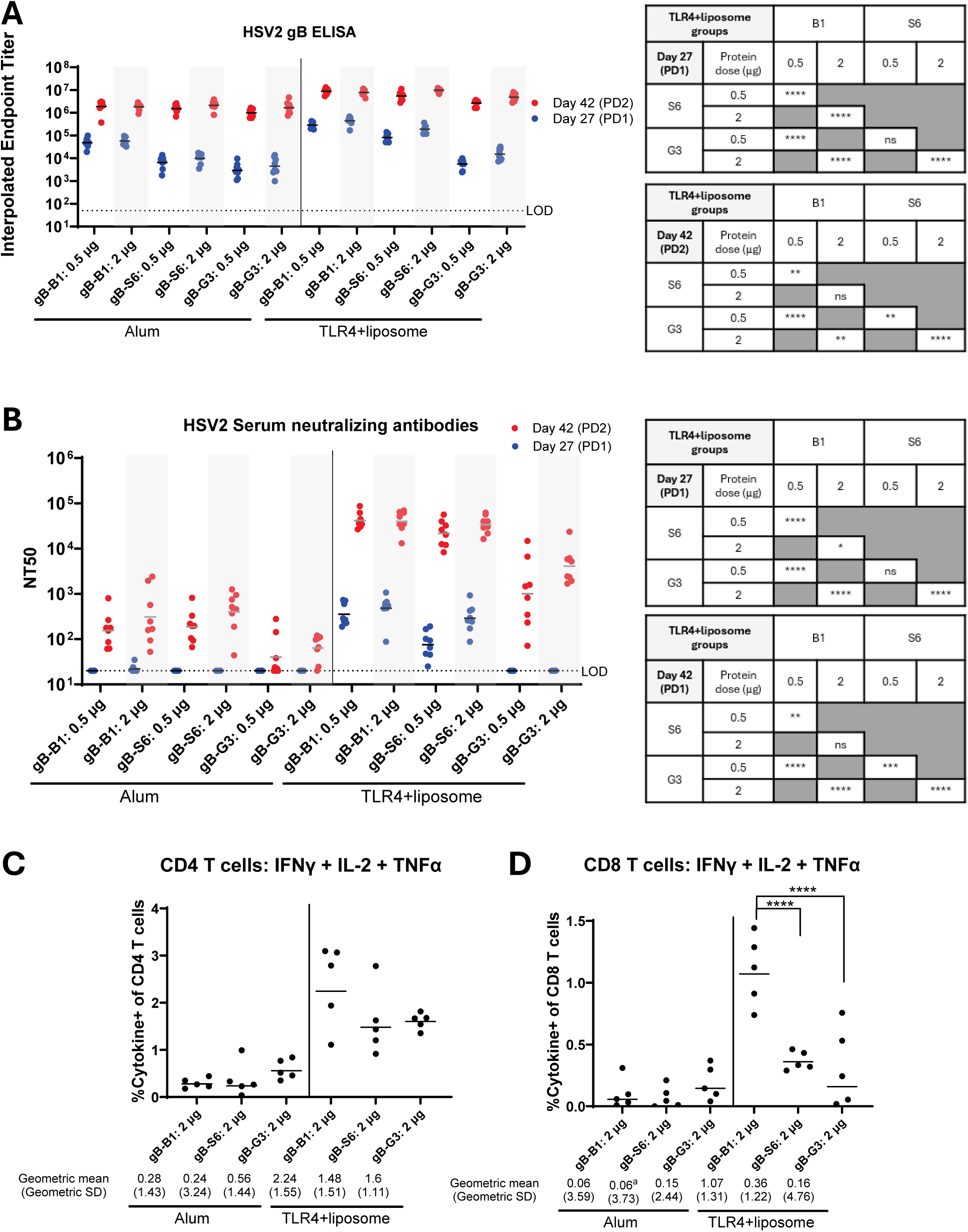
Antibody and T cell responses from protein immunizations. Antibody responses are characterized by **(A)** ELISA to quantify HSV-2 gB binding titers or **(B)** neutralization assay to quantity HSV-2 serum neutralizing antibody titers, where symbols represent individual animals (n=8) and bars represent geometric mean. Statistics are shown in tables and were conducted by two-way ANOVA with multiple comparisons for all groups, and statistically significant comparisons in TLR4+liposome groups reported in the tables where ns = not significant, *p<0.05, **p<0.01 and ****p<.0001. T cell responses in **(C)** CD4 and **(D)** CD8 T cell populations were characterized by T cell intracellular cytokine staining (ICS) from splenocytes isolated at day 42 (2 weeks post-dose 2). The sum total frequency of T cells producing any Th1 cytokine (IFNγ, IL-2, TNFα) for individual animals (n= 5) from the 2 μg protein dose groups are shown by symbols, with bars representing geometric mean. Statistical analysis was conducted using a two-way ANOVA with multiple comparisons where ****p<0.001. Values for geometric means and geometric standard deviation (SD) are calculated from 5 individual animals unless otherwise indicated. ^a^Values for geometric means and geometric SD are calculated from 4 individual animals as one animal had undetectable levels of cytokine-producing T cells.

To further characterize the antibody responses, we quantified the titers of serum neutralizing antibodies from immunized mice. Neutralizing antibody titers reached higher magnitude in TLR4+liposome groups compared to alum adjuvant groups, with little to no detectable neutralizing antibody induced at day 27 using alum as an adjuvant. However, as observed in quantification of the ELISA titers, neutralizing antibody titers were boosted in all adjuvant groups after the second immunization, and statistically significant differences in GMTs were only detected in immunization groups using TLR4+liposome as an adjuvant (**Figure 5B**). Similar to trends observed in the ELISA, gB-B1 immunizations induced higher neutralizing antibody titers compared to gB-S6 and gB-G3 groups that reached statistical significance in TLR4+liposome adjuvant groups at days 27 and 42 (**Figure 5B**). As observed with ELISA GMTs, gB-B1 induced a higher fold change at day 42 in neutralizing titers versus gB-G3 vaccines (3.9 to 41.4-fold) and compared to gB-S6 vaccines (0.8 to 1.9-fold) (**Supplementary Table 2**). Immunization with gB-S6 also induced higher serum neutralization titers compared to gB-G3 at day 42 (range: 4.7 to 21.4-fold; **Supplementary Table 2**), with gB-G3 immunization being unable to induce detectable serum neutralization titers irrespective of adjuvant use or antigen dose at day 27 (**Figure 5B**). Together, these data suggest that gB-B1 immunizations generally induced more robust antibody responses compared to gB-S6 and gB-G3 protein antigens, with gB-G3 being the least immunogenic of the three constructs.

To ensure that detection of gB-binding antibodies was not biased to prefusion or postfusion conformations of HSV-2 gB protein, we quantified serum antibody titers using gB-B1 (postfusion) or gB-G3 (prefusion) proteins as the coating antigen by ELISA. The magnitude and pattern for antibody binding titers were nearly identical between assays utilizing gB-B1 and gB-G3 proteins as coating antigens (**Supplementary Figure S5**), indicating that the overall trends regarding the relative immunogenicity of gB-B1, gB-S6 and gB-G3 protein were not dependent on the conformation of the coating antigen used in the ELISA.

We also examined splenic CD4 and CD8 T cell responses in protein-immunized mice by peptide pool stimulation and intracellular cytokine staining for Th1 cytokine production. Alum immunization induced very weak or non-detectable cytokine production in CD4 and CD8 T cells, irrespective of the antigen used in immunizations (**Figure 5C,D**). In contrast, immunization groups using TLR4+liposome induced robust CD4 and CD8 T cell cytokine production for all antigen groups. Similar to the trends observed for antibody responses, gB-B1 protein immunization with TLR4+liposome induced the highest levels of cytokine production in CD4 and CD8 T cells compared to gB-S6 and gB-G3 groups (**Figure 5C,D**). This effect was more pronounced in CD8 T cells compared to CD4 T cells and reached statistical significance (**Figure 5D**).

In summary, these data demonstrate that immunizations with postfusion HSV-2 gB-B1 protein were more immunogenic compared to prefusion HSV-2 gB constructs gB-S6 and gB-G3. Moreover, gB-S6 protein induced higher antibody titers compared to gB-G3 protein immunization but showed no advantages in ability to induce CD4 or CD8 T cell Th1 cytokine production.

### Immunogenicity of mRNA encoding prefusion and postfusion HSV-2 gB in mice

To further evaluate the immunogenicity of our prefusion and postfusion HSV-2 gB constructs, we generated mRNA sequences for the ectodomains of HSV-2 gB-B1, gB-S6, and gB-G3 designs encapsulated into a lipid nanoparticle (LNP) and immunized mice. mRNA immunization groups utilized a low dose (0.3 µg total mRNA) or high dose (1.2 µg total mRNA), with protein controls using 2 µg of gB-B1 or gB-S6 protein with TLR4+liposome adjuvant. mRNA immunizations induced robust HSV-2 gB binding antibody titers in all groups and timepoints examined. There were no statistically significant differences in GMTs between gB-B1 mRNA immunizations compared to gB-S6 or gB-G3 mRNA immunizations at day 27 or 42 at the 0.3 µg dose (**Figure 6A, Supplementary Table 3**). There were also no differences in GMTs between gB-S6 and gB-G3 mRNA immunizations at 0.3 µg at any time point. At the 1.2 µg dose, gB-B1 mRNA also showed no statistically significant differences in GMTs compared to immunization with gB-S6 or gB-G3 mRNA at day 27 (**Figure 6A, Supplementary Table 3**). At day 42, there were statistically significant increases in GMTs between gB-B1 mRNA and gB-S6 or gB-G3 mRNA immunization, but this only amounted to a 1.7 and 2.2-fold change, respectively (**Supplementary Table 3**). As observed in the protein immunization experiment, there were no differences in the overall trends in HSV-2 gB-specific antibody titers irrespective of gB-B1 or gB-G3 proteins being used as coating antigens in the ELISA, thus indicating that the relative immunogenicity of gB-B1, gB-S6, and gB-G3 mRNA constructs was not biased toward coating antigen conformation (**Supplementary Figure S6**).

**Figure 6.**
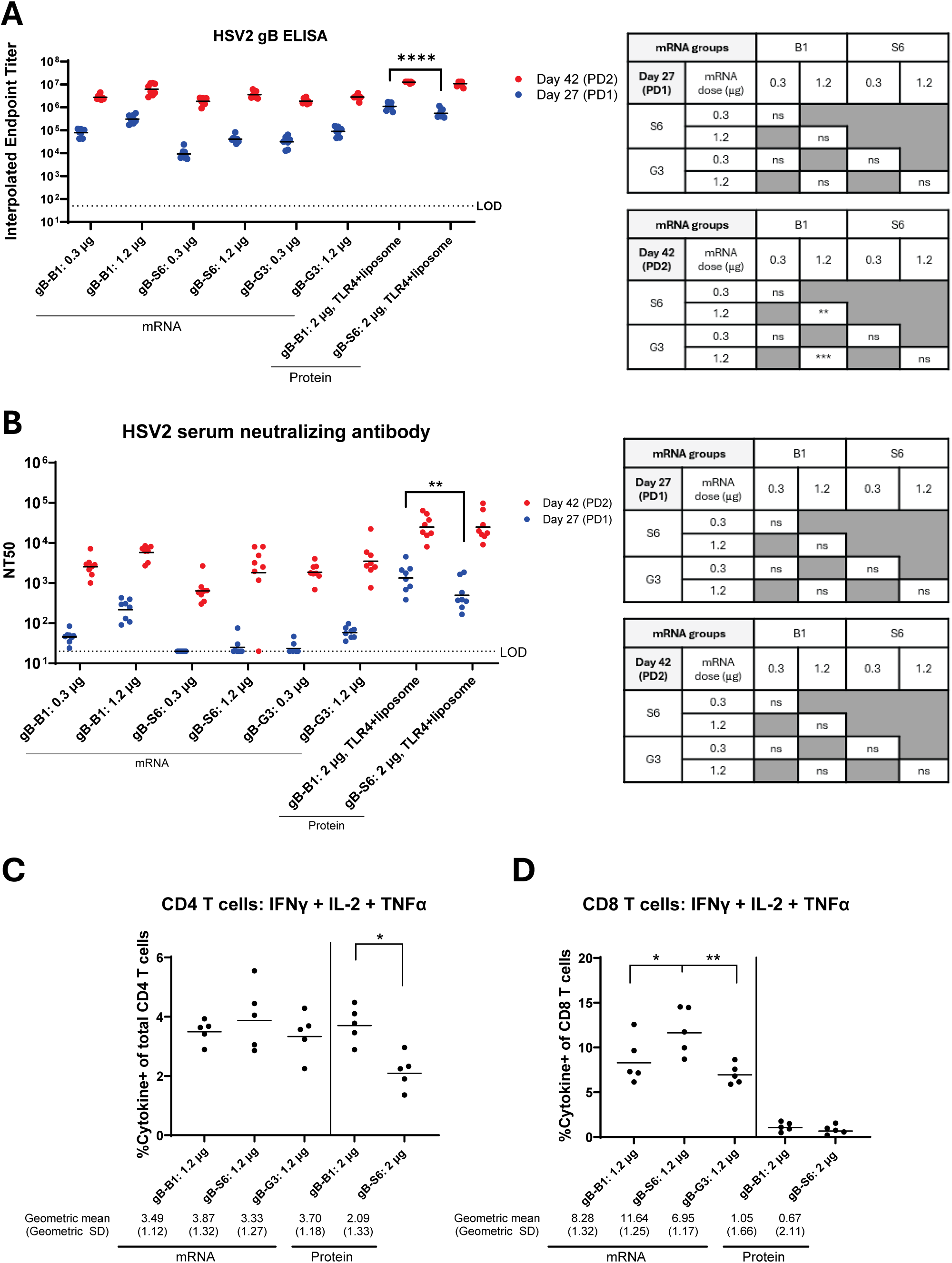
Antibody and T cell responses from mRNA immunizations. Antibody responses are characterized by **(A)** ELISA to quantify HSV-2 gB binding titers or **(B)** neutralization assay to quantify HSV-2 serum neutralizing antibody titers, where symbols represent individual animals (n=8) and bars represent geometric mean. Statistics are shown in tables and were conducted by one-way ANOVA with multiple comparisons for all groups, and statistically significant comparisons in mRNA groups reported in the tables and protein groups reported on the graphs where ns = not significant, *p<0.05, **p<0.01 and ***p<.001. T cell responses in **(C)** CD4 and **(D)** CD8 T cell subsets were characterized by T cell intracellular cytokine staining (ICS) from splenocytes isolated at day 42 (2 week post-dose 2). The sum total frequency of T cells producing any Th1 cytokine (IFNγ, IL-2, TNFα) for individual animals (n=5) from the 1.2 μg mRNA or 2 μg protein groups are shown by symbols, with bars representing geometric mean. Statistical analysis was conducted using a two-way ANOVA with multiple comparisons where *p<0.05 and **p<0.01. Values for geometric means and geometric standard deviation (SD) are calculated from 5 individual animals.

mRNA immunization induced weak or non-detectable serum neutralizing antibody responses at day 27, and robust responses were only detected after the second immunization (**Figure 6B**). This contrasts with protein immunization, which induced robust serum neutralization titers as early as day 27 (**Figure 5B**). Interestingly, there were no statistically significant differences in the magnitude of serum neutralizing antibody GMTs between the gB-B1, gB-S6, and gB-G3 mRNA immunization groups in any dose-matched comparisons at days 27 or 42 (**Figure 6B**). *In vitro* cell-based assays using HiBiT and non-HiBiT tagged mRNA constructs confirmed equivalent protein expression between gB-B1, gB-S6, and gB-G3 ectodomain constructs (**Supplementary Figure S7**), suggesting equivalent antigen expression corresponding to comparable immunogenicity between these constructs *in vivo*. In addition, immunization with 2 µg gB-B1 protein with TLR4+liposome recapitulated results from earlier studies (**Figure 5B**) showing increased neutralizing antibody titers compared to 2 µg gB-S6 protein with TLR4+liposome immunizations at day 27 (**Figure 6B, Supplementary Table 3**), thus confirming previous findings (**Figure 5B**). In summary, these data demonstrate that mRNA vaccination with postfusion gB-B1 and prefusion gB-S6 and gB-G3 constructs induced comparable antibody responses *in vivo*.

mRNA immunization also induced robust CD4 and CD8 T cell responses (**Figure 6C, D**). CD4 responses were comparable among gB-B1, gB-S6, and gB-G3 mRNA constructs (**Figure 6C**). gB-S6 mRNA immunization induced increased CD8 T cell responses, reaching statistical significance when compared to gB-B1 and gB-G3 mRNA groups (1.4- and 1.7-fold, respectively (**Figure 6D**). There were no statistically significant differences in CD8 T cell responses induced by gB-B1 versus gB-S6 protein immunization (**Figure 6D**), which contrasts with the prior protein immunization study (**Figure 5D**).

In summary, although we observed a statistically significant increase in binding antibody, serum neutralization titers, and CD8 T cells induced by gB-B1 postfusion versus gB-S6 and gB-G3 prefuson antigens in the protein immunization experiment (**Figure 5**), these results were not recapitulated when these constructs were expressed using mRNA (**Figure 6**). Together, this suggests that prefusion HSV-2 gB constructs do not elicit superior immune responses in mice when compared to postfusion HSV-2 gB.

## Discussion

HSV gB is responsible for fusing the viral envelope with the host plasma membrane (*30, 31*). The protein is highly conserved across HSV strains and shows substantial sequence homology between HSV-1 and HSV-2 (*32*). During natural infection, gB elicits robust antibody responses that can contribute significantly to virus neutralization (*33, 34*). Given the central role of gB in viral entry and its significance as a vaccine antigen, elucidating its native, immunologically relevant prefusion structure will not only advance our understanding of the molecular mechanism underlying HSV membrane fusion but also facilitate structure-guided vaccine design.

Here, we identified a set of prefusion-stabilizing substitutions for HSV-2 gB that are similar to those previously identified for HCMV gB (*16*). These substitutions successfully stabilized HSV-2 gB-G3 in the prefusion conformation, revealing additional structural details that appear to be unique to HSV gBs (**Figure 4**). Our protein engineering work on HSV-2 gB further underscores the importance of interprotomer disulfide substitutions in stabilizing gB in a prefusion-like conformation. This strategy was also recently applied to stabilize EBV gB in the prefusion conformation (*17*). McCool et al. found that for both B cells and epithelial cells, immunization with prefusion-stabilized EBV gB resulted in modestly improved neutralizing titers relative to immunization with postfusion EBV gB, and that depletion of human sera from EBV-positive donors by prefusion EBV gB resulted in a greater reduction in the IC₅₀ compared to depletion with postfusion gB. With these findings, we had reason to suspect that a fully prefusion-stabilized HSV-2 gB construct may elicit an enhanced immune response by displaying prefusion-specific epitopes that can be targeted by the immune system to prevent viral fusion.

Immunization with HSV-2 gB protein demonstrated that postfusion gB-B1 induced higher binding and serum neutralization titers as well as CD8 T cell responses compared to prefusion -S6 and -G3 constructs (**Figure 5)**. These trends were not recapitulated when the vaccination was performed with mRNA (**Figure 6)**. It is possible that inherent differences affecting antigen delivery, antigen persistence, and protein properties between these two vaccine modalities accounts for these results. Indeed, these differences between immunogenicity for mRNA and recombinant protein vaccination have been previously observed in other vaccines tested in rodents (*35–37*).

Regardless of mode of antigen delivery, our studies did not demonstrate superior immunogenicity of prefusion HSV-2 gB-S6 or gB-G3 constructs for immunization compared to postfusion gB-B1 constructs in mice. Recent work with HSV-1 gB has identified a prefusion-specific nanobody targeting a neutralizing epitope within DIV, DI, and DIII (*28*). Specifically, the nanobody bound to HSV-1 and HSV-2 gB, which suggests the presence of conserved neutralizing epitopes in prefusion HSV-2 gB. Another prefusion-specific antibody was recently isolated from mice immunized with a prefusion-stabilized HSV-1 gB construct, and that antibody binds an epitope at the interface between DI and DII (*18*). It remains to be determined whether humans elicit similar prefusion-specific gB-directed antibodies.

Considering ongoing antibody isolation efforts, neutralization-sensitive epitopes specific to the prefusion conformation of gB appear to be rare, further suggesting that additional strategies may be required to develop an effective gB-based HSV-2 vaccine, including the display of multiple antigens, multimerization, and immune focusing. Stabilization of gB in the prefusion conformation, which is the only conformation expected to interact with gH/gL (*38*), may enable the formation of the gB/gH/gL complex, which may be an effective candidate vaccine antigen. Overall, the findings presented here are in agreement with recent publications, showing that maintaining the prefusion conformation of herpesvirus fusion proteins is not sufficient to elicit a significantly superior immune response relative to the postfusion conformation (*16–18, 39*). Our development of a stabilized ectodomain of HSV-2 gB in its prefusion conformation will assist these efforts, allowing for the evaluation of conformationally specific interactions with other viral surface glycoproteins and the isolation of prefusion-specific antibodies, which may be valuable in the development of effective monoclonal antibody therapeutics for the treatment of HSV-2 infections.

## Acknowledgements

We thank members of the McLellan Laboratory for providing helpful comments on the manuscript and Dr. Axel Brilot and Dr. Evan Schwartz at the Sauer Structural Biology Laboratory at UT Austin for assistance with cryo-EM data collection. We thank Marco Blanco for formulating the LNP particles and Dr. Emmanuel Ndashimye for characterizing the expression of the gB mRNAs. We thank Uijin Jeong and Silveria Rodriguez for performing the ELISA and neutralization assays. We acknowledge the University of Texas College of Natural Sciences and award RR160023 of the Cancer Prevention and Research Institute of Texas for support of the EM facility at the University of Texas at Austin. This work was funded by Merck & Co., Inc., Rahway, NJ, USA.

## Author Contributions

Conceptualization – M.R.S., P.O.B., L.Z., E.D., D.W. and J.S.M.; Investigation and visualization – M.R.S., P.O.B., Y.S., C.W., M.D.S., N.V.J., W.A.R., and J.S.M.; Writing – Original Draft, M.R.S., D.M., and J.S.M.; Writing – Reviewing & Editing – M.R.S., D.M., P.O.B., C.W., N.T., M.D.S., N.V.J., L.Z., E.D., D.W., and J.S.M.; Supervision, E.D., D.W., and J.S.M.

## Materials and Methods

### Design scheme for prefusion-stabilized HSV-2 gB variants

The initial HSV-2 gB base construct (gB-B0) comprises residues 1–731 of the ectodomain from the HG52 strain (UniProtKB: P08666), followed by a C-terminal T4 fibritin trimerization domain (foldon), a human rhinovirus 3C (HRV3C) protease recognition site, an octa-histidine tag, and a Twin-Strep tag. The gene encoding this construct was cloned into the pαH mammalian expression vector and sequence-verified. A modified base construct (gB-B1) was generated by deleting the proline-rich region (residues 26–73) following the signal peptide. All further HSV-2 gB variants were based on this plasmid. A homology model of the HSV-2 gB ectodomain in the prefusion conformation was built using SWISS-MODEL (*24*) and a previously reported detergent-solubilized structure of full-length HCMV gB (PDB ID: 7KDP) (*25*), rebuilt and relaxed using ModelAngelo (*40*) and Isolde (*41*). Structure-based design of disulfide bonds, polar interactions, cavity-filling substitutions, and proline substitutions was guided by the HSV-2 model and sequence alignments with gB homologs from other herpesviruses. Combinatorial variants were generated to assess whether additive stabilization effects could be achieved.

### Purification of HSV-2 gB variants by affinity and size-exclusion chromatography

Plasmids encoding HSV-2 gB variants were transiently transfected into 40 mL cultures of FreeStyle 293-F cells using 25 kDa linear polyethyleneimine (PEI, Polysciences). After incubation at 37 °C with 8% CO₂ for 5–7 days, culture supernatants were harvested by centrifugation and sterile-filtered through 0.22 µm membranes. Clarified supernatants were applied to Strep-Tactin XT 4Flow high-capacity resin (IBA Lifesciences) by gravity flow. Following loading, the resin was washed with 3 column volumes of phosphate-buffered saline (PBS) and protein was eluted using BXT elution buffer (100 mM Tris-HCl pH 8.0, 150 mM NaCl, 1 mM EDTA, 50 mM biotin; IBA Lifesciences). Eluted fractions were assessed by SDS-PAGE, and those containing HSV-2 gB were pooled and concentrated using Amicon Ultra centrifugal filters (MilliporeSigma). Concentrated protein was flash-frozen in liquid nitrogen and stored at −80 °C. For size-exclusion chromatography (SEC), samples were thawed at room temperature and injected onto a Superose 6 Increase 10/300 GL column (Cytiva) pre-equilibrated in 20 mM Tris-HCl pH 8.0, 200 mM NaCl, and 0.02% (w/v) sodium azide. Eluted fractions corresponding to the major peak were pooled, concentrated, aliquoted, and flash-frozen in liquid nitrogen for further analysis.

### Expression and purification of Fabs for structural and biophysical studies

A stop codon was introduced before the hinge region of the heavy chains to generate fragments of antigen binding (Fabs) for 2c, D48, HSV010-13, and BMPC-23. FreeStyle 293-F cells were transiently transfected with plasmids encoding the corresponding heavy and light chains using PEI. Cultures were incubated at 37 °C with 8% CO₂ and harvested six days post-transfection. Supernatants were clarified by centrifugation, filtered through 0.22 μm membranes, and concentrated using a tangential flow filtration cassette (Pall). Fabs were purified using CaptureSelect IgG-CH1 Affinity Matrix (ThermoFisher), eluted with 100 mM glycine pH 3.0, and immediately neutralized with 1 M Tris-HCl pH 8.0. Fab-containing eluates were further purified by SEC on a HiLoad 16/600 Superdex 200 pg column (Cytiva) equilibrated in 20 mM Tris-HCl pH 8.0, 200 mM NaCl, and 0.02% (w/v) sodium azide. Final protein preparations were concentrated to 1–10 mg/mL using Amicon Ultra centrifugal filters (MilliporeSigma), flash-frozen in liquid nitrogen, and stored at −80°C.

### Differential scanning fluorimetry

Reactions were prepared in white-walled 96-well PCR plates (VWR) with 0.45 mg/mL protein and 15X SYPRO Orange (ThermoFisher; 5,000X stock in DMSO) in a total volume of 50 µL per well. The absorbance of the working SYPRO Orange solution was 0.295 at 468 nm, measured using a NanoDrop One spectrophotometer (ThermoFisher) with a 1 mm path length. Plates were sealed and placed in a LightCycler 480 II instrument (Roche) equipped with a 100 W xenon excitation lamp. Fluorescence was monitored using a 465 nm excitation filter (25 nm half-bandwidth) and a 580 nm emission filter (20 nm half-bandwidth) while the temperature was increased from 25 to 90 °C. Fluorescence data were plotted as the first derivative of fluorescence intensity with respect to temperature (dF/dT) to determine thermal transition midpoints (Tm).

### Negative-stain electron microscopy

Purified and concentrated HSV-2 gB-B1 protein was incubated with Fabs 2c, HSV010-13, and BMPC-23 at a 2:1 molar ratio (Fab:gB) for 15–30 minutes at room temperature. The complex was diluted to a final concentration of 0.04 mg/mL in SEC buffer and immediately applied to glow-discharged 400-mesh copper grids coated with carbon-stabilized Formvar film (Electron Microscopy Sciences). Grids were stained with 2% (w/v) methylamine tungstate (Nano-W; Nanoprobes) and imaged using a JEOL NEOARM transmission electron microscope operated at 200 kV. Micrographs were acquired at a nominal magnification of 50,000x (corresponding to a pixel size of 2.16 Å) using a OneView CMOS camera (Gatan) in 4k × 4k mode with Digital Micrograph software (Gatan). Images were imported into cryoSPARC (Structura Biotechnology) for contrast transfer function (CTF) estimation, particle picking, and two-dimensional classification (*42*).

### Cryo-EM sample preparation and data collection

Cryo-EM grids (CF-400 1.2/1.3; Electron Microscopy Sciences) were glow-discharged for 30 s at 15 mA using a PELCO easiGlow system prior to sample application. Samples were prepared in EM buffer containing 2 mM Tris-HCl pH 8.0, 200 mM NaCl, 0.02% (w/v) sodium azide, 3% (v/v) glycerol, and 0.01% (w/v) amphipol A8-35.

For the gB-S2 complex, gB-S2 was incubated with a 1.5-fold molar excess of 2c Fab and a 1.2-fold molar excess of D48 Fab in EM buffer for 30 min at 4 °C, yielding a final concentration of 2.1 mg/mL. Immediately before vitrification, CHAPS (VitroEase Buffer Screening Kit; ThermoFisher) was added to a final concentration of 0.1X CMC. A 3 µL aliquot of the complex was applied to each grid, double-blotted, and plunge-frozen in liquid ethane using a Vitrobot Mark IV (ThermoFisher) operated at 100% humidity and 4 °C with a blot time of 3 s, blot force of −1, and wait time of 5 s.

For the gB-G3 complex, gB-G3 was mixed with 1.2-fold molar excesses of 2c and D48 Fabs to a final concentration of 3.5 mg/mL, incubated for 30 min at 4 °C, and supplemented with CHAPS to a final concentration of 0.2X CMC. Three microliters of the complex solution were applied to glow-discharged grids, double-blotted, and plunge-frozen under identical conditions except with a blot time of 5 s and blot force of 6.

Cryo-EM data were collected on an FEI Titan Krios transmission electron microscope (ThermoFisher) operated at 300 kV and equipped with a K3 direct electron detector (Gatan). Micrographs were recorded at a nominal magnification of 105,000x, corresponding to a calibrated pixel size of 0.83 Å. A total of 17,634 micrographs were collected for the gB-S2 complex, all at a 30° tilt. For the gB-G3 complex, a total of 11,000 exposures were collected, 5,182 with a 30° tilt and 5,818 without tilt. Data acquisition was performed using SerialEM v4.1 (*43*).

### Cryo-EM data processing, model building, and refinement

Patch-based motion correction, patch-based CTF estimation, micrograph curation, particle picking, and 2D classification were initially performed in cryoSPARC Live (*42*). Selected micrographs and particles were then exported to cryoSPARC v4.5 for additional 2D classification, *ab initio* reconstruction, heterogeneous refinement, homogeneous refinement, and non-uniform refinement of final classes.

Initial models were built with ModelAngelo (*40*), and iterative model building and refinement of the gB-G3 structure were performed in Phenix (*44*), Coot (*45*), and Isolde (*41*). Buried surface areas and interacting residues were calculated using the Protein Interfaces, Surfaces and Assemblies (PISA) server (*46*) at the European Bioinformatics Institute. Structural figures were prepared in ChimeraX (*47*). Data collection and refinement statistics are presented in **Supplementary Table 1**.

### Vaccine formulations

Protein vaccines were formulated for final antigen dose at 0.5 or 2 µg with either 25 µg of alum or 2.5 µg TLR4+liposome adjuvant. Linear mRNA constructs encoding the ectodomains of gB-B1, gB-S6, and gB-G3 were produced at Merck & Co., Inc. (West Point, PA). mRNA coding sequences were codon-optimized using Merck & Co., Inc., Rahway, NJ, USA algorithm. Plasmid templates of gB-B1, gB-S6, and gB-G3 were synthesized by vendor (GenScript, USA) and linearized for *in vitro* transcription (IVT). Following IVT, the produced mRNAs were purified by oligo(dT) affinity chromatography (Sartorius). Expression of all three mRNAs was confirmed in cell-based assays (**Supplementary Figure S7**). Linear mRNA vaccines were formulated for final RNA dose at 0.5 or 1.2 µg in lipid nanoparticles (LNP). LNP were prepared and characterized as previously described (*48*). The LNPs are comprised of an ionizable amino lipid ((13Z,16Z)-N,N-dimethyl-3-nonyldocosa-13,16-dien-1-amine), distearoylphosphatidylcholine (DSPC), cholesterol, and poly(ethylene glycol)2000-dimyristoylglycerol (PEG2000-DMG), in a molar ratio of 58:30:10:2, respectively.

Lipid nanoparticle size was determined by dynamic light scattering using a DynaPro particle sizer (Wyatt Technology, Santa Barbara, CA) and measured at 80–100 nm across all formulations. The concentrations of the four lipids in LNP were determined by gradient reverse-phase ultrahigh-performance liquid chromatography methods. Additionally, encapsulation efficiency of RNA, as measured by the SYBR gold fluorimetric method, was greater than 95% for all LNP preparations tested.

### Mouse immunizations

We used 9-week-old female BALB/c mice purchased from Charles River Laboratories. Food and water were provided ad libitum, with a 12-hour light/dark cycle. All animal experiments were approved by the Institutional Animal Care and Use Committee (IACUC), Merck & Co., Inc. (Rahway, NJ, USA). Mice were immunized with two doses of vaccine intramuscularly at week 0 and week 4. At days 27 (∼4 weeks post-dose 1, PD1) and 42 (2 weeks post-dose 2, PD2), mice were bled and sera samples collected for antibody analyses. Spleens were also harvested at day 42 for T cell analysis.

### Antibody ELISA

HSV-2 gB antibody binding titers were assessed using an enzyme-linked immunosorbent assay (ELISA). Briefly, HSV-2 gB protein coating antigen at a concentration of 1 µg/mL was captured onto 384-well MaxiSorp assay plates (Thermo Fisher, catalog #460518) overnight at 4 °C. The next day, plates are washed and blocked with 3% non-fat dry milk in PBS with 0.05% TWEEN-20 (PBST) before diluted sera addition. Serially diluted sera (50-fold initial dilution and 4-fold, 10-point titration) or mouse anti-HSV-2 gB monoclonal antibodies as positive controls were added to blocked ELISA plates for 2 hours at 22 °C. Goat anti-mouse IgG conjugated with horseradish peroxidase was subsequently added at a concentration of 0.25 µg/mL and incubated for 1 hour at 22 °C before developing with a chemiluminescent substrate (equal parts Luminol/enhancer solution and stable peroxide solution). Luminescence was read using the EnVision multimode reader (Perkin-Elmer). Interpolated endpoint titers were determined with the Interpolation formula for titer based on data at dilutions above and below designated cut-off (ie. 50,000). The interpolated endpoint titers were graphed by treatment group as the geometric mean + 95% confidence interval in Prism (GraphPad Software).

titer = (starting dilution / series dilution factor) x (series dilution factor^t)

where t = x - [(cut-off - L) / (H - L)]

H = High well OD = the OD of the first well ABOVE the cutoff

L = Low well OD = the OD of the first well BELOW the cutoff

x = Low well number = the column # of the "low well OD"

cut-off = OD of designated cut-off, such as 50,000

### HSV-2 serum neutralization assay

Serum neutralizing antibody titers were assessed using a serum neutralization assay with an HSV-2-GFP reporter virus (strain MS, produced by Dr. Hua Zhu’s laboratory, Rutgers University, New Jersey) and baby hamster kidney-21 (BHK-21) cells (ATCC, catalog # CCL-10). Mouse sera samples were heat-inactivated for 30 minutes at 56 °C before use. Sera samples were diluted 1:20 in serum-free Eagle’s Minimum Essential Media (EMEM) with culture media supplements and 5% (v/v) baby rabbit complement (Cedarlane, catalog #CL3441). The diluted serum samples as well as a mouse-anti HSV gD protein monoclonal antibody (Native Antigen Company, clone E317, catalog #MAB12121-200) as positive controls were serially diluted 3-fold. HSV-2-GFP reporter virus was added at 1.0 x 10^4^ plaque-forming units to the sera samples and the control antibody in equal volume. After one hour incubation at 37 °C, 4 x 10^3^ BHK-21 cells per well were added to the 384-well plates. Virus neutralization proceeded for 18–20 hours at 37 °C until quantification of GFP-positive foci using an Acumen (SPT Labtech) high-throughput imager. To determine half-maximal neutralization titers (NT_50_), GFP fluorescent signal in each well was converted to a percentage of infectivity by dividing E*_max_* (infected cells) by E*_min_* (uninfected cells). This percentage for each sample dilution series was fit to a 4-parameter logistic curve (PerkinElmer version 12.1.10.1) and graphed in Prism (GraphPad Software).

### Murine splenocyte peptide stimulation

T cell Th1 cytokine production was assessed in murine splenocytes at day 42 after immunization. Spleens from individual mice were processed to generate single-cell splenocyte suspensions. After red blood cell lysis, 1–2 x 10^6^ cells were plated into round-bottom 96-well plates and stimulated with 2 µg/mL of peptide pools spanning the HSV-2 gB protein (15 mers with 11 amino acid overlap; JPT Peptide Technologies GmbH, Berlin, Germany) or no-peptide negative controls (0.1% dimethyl sulfoxide v/v, Sigma-Aldrich catalog #D2650) in the presence of 1.25 μg/mL soluble anti-mouse CD28 (clone 37.51, BD Biosciences catalog #557393) and 1.25 μg/mL soluble anti-mouse CD49d (clone R1-2, BD Biosciences catalog #553154) for 6 hours total at 37 °C, with the addition of Brefeldin A (Sigma-Aldrich catalog #B7651) for the final 5 hours at a concentration of 10 µg/mL. Ethylenediaminetetraacetic acid (EDTA) was used to arrest the reaction, and cells were stored overnight at 4 °C prior to immunostaining.

### T cell intracellular cytokine staining (ICS) and flow cytometry

Stimulated splenocytes were immunostained for surface markers to identify T cell populations, followed by ICS to identify Th1 cytokines tumor necrosis factor α (TNFα), interleukin-2 (IL-2) and interferon γ (IFNγ). The following antibodies and stains were utilized: LIVE/DEAD Fixable Violet Dead Cell Stain (Invitrogen catalog #L34955); CD3-APC Cy7 (Clone 145-2C11, BD Biosciences catalog #557596); CD4-PE (Clone RM4-5, BD Biosciences catalog #553049); CD8α-BUV395 (Clone 53-6.7, BD Biosciences catalog #563786); TNFα-PE Cy7 (Clone MP6-XT22, BD Biosciences catalog #557644); IL-2-PCF594 (Clone JES6-5H4, BD Biosciences catalog #562483); IFNγ-APC (Clone XMG1.2, BD Biosciences, catalog #554413). Surface and intracellular staining was conducted in the presence of Mouse Fc Block (Clone 2.4G2, BD Biosciences #553142). ICS was conducted according to manufacturer’s instructions using Cytofix/Cytoperm buffer (BD Biosciences catalog #554722) and Perm/Wash Buffer (BD Biosciences catalog #554723), and cells fixed using BD Stabilizing Fixative (BD Biosciences catalog #338036) prior to flow cytometric analysis on a Fortessa X-50 flow cytometer (BD Biosciences). Data was analyzed using FlowJo^TM^ software (BD Biosciences, version 10.8.1), and graphed in Prism (GraphPad software).

### Epitope structural analysis

To structurally compare the potential epitopes present on the prefusion (G3) and postfusion HSV-2 gB structures, we first downloaded a model of the postfusion conformation from the Protein Data Bank (PDB ID: 8RH1) and removed the bound Fab. Chain IDs and residue numbers were altered to be consistent between the prefusion and postfusion conformations. In each conformation, per-residue solvent accessible surface area (SASA) values were calculated using the Shrake Rupley algorithm implemented within the BioPython SASA module (*49*). Per-residue SASA values in each conformation were normalized to the maximum experimentally determined value for that residue type (*50*), leading to a relative scale (rSASA) from 0 (fully buried) to 1 (fully accessible) in each conformation. To compare across the two conformations, per-residue rSASA values were subtracted (prefusion – postfusion). In this metric, a score of 1 indicates a residue that is fully accessible in the prefusion conformation and fully buried in the postfusion conformation, a value of –1 indicates a residue that is fully accessible in the postfusion conformation and fully buried in the prefusion conformation, and a value of 0 indicates that the residue is equally accessible in both conformations. In addition to rSASA analysis, for each residue in each conformation, we measured all other residues within 10 Å (Cα-Cα) and defined this list as that residue’s neighborhood using Biopandas (*51*). For each residue we compared the similarity among the neighborhood lists in each conformation via Jaccard similarity metric, where a value of 1 indicates that the lists are identical (neighborhood is the same in both conformations) and a value of 0 indicates that there are no similarities between the lists. rSASA values and neighborhood Jaccard similarities between the two conformations were assigned to the B-factor column of the PDB files and visualized in PyMOL software via the spectrum command.

### HiBiT Assay in 293F cells

HSV-2 gB variants B1, S6, and G3 mRNAs were synthesized by IVT and purified by Monarch® Spin RNA Cleanup Kit (500 μg) (NEB, catalog# T2050). To facilitate expression profiling, an 11-amino acid HiBiT tag (Promega) was fused to each gB construct. The HiBiT tag complements its partner protein, LgBiT, to reconstitute an active luciferase enzyme. In the presence of substrate, the resulting bioluminescence signal quantitatively reflects the expression level of the tagged protein. For transfection, 50 ng of HiBiT-tagged B1, S6, or G3 mRNA was formulated with Lipofectamine MessengerMax (Invitrogen, catalog# LMRNA150) according to the manufacturer’s instructions. The mRNA-Lipofectamine complexes were serially diluted two-fold across 10 dilution points and used to transfect 293 cells. Bioluminescence readings were recorded hourly for up to 48 hours post-transfection to profile expression kinetics (Promega, catalog# N2572). Expression levels for each construct were plotted as dose-response curves over time. The area under the curve (AUC) was calculated to quantify overall expression. To specifically assess intracellular expression, 293 cells stably expressing LgBiT endogenously were used, allowing luminescence signals to represent intracellular HiBiT-tagged protein levels. For extracellular expression analysis, parental 293 cells were employed; exogenous LgBiT protein (Promega, catalog# N401A) was added to the culture medium to enable luminescence detection exclusively from secreted HiBiT-tagged proteins.

### Western Blot in 293F cells

Protein expression was analyzed bycapillary-based immunoassay using Jess Automated Western Blot System (ProteinSimple, USA). Briefly, FreeStyle 293-F cells were lysed 24 hr post-transfection using Pierce IP Lysis buffer (ThermoFisher, cat #87787) and protease inhibitor cocktail (ThermoFisher, cat # 78442). BCA assay was used to measure protein concentration (ThermoFisher, cat # 23227). Total protein was detected using total protein detection module (ProteinSimple, USA, cat # DM-TP01) and HSV-2 gB proteins were probed using primary antibody produced at Merck & Co., Inc. (West Point, PA) and anti-mouse detection module (ProteinSimple, USA, cat # DM-002). Samples were separated using capillary cartridge (ProteinSimple, USA, cat # SM-W004) and detection of both HSV-2 gB and total protein in a single run was enabled using Replex module (ProteinSimple, USA, cat # DM-002). Data was analyzed on Compass for Simple Western software (Bio-Techne, USA).

