## Supplemental Figures and Tables for "Structure-based design of soluble prefusion-stabilized herpes simplex virus type 2 glycoprotein B antigens"

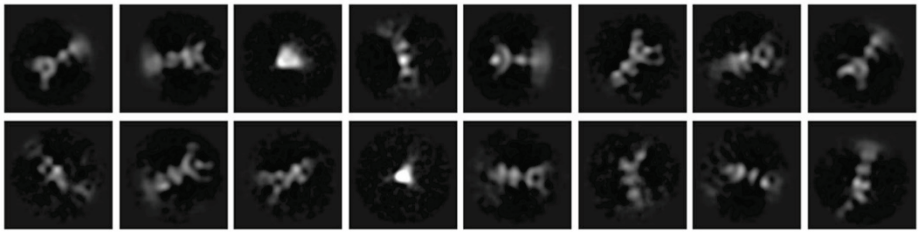

**Supplementary Figure S1: Negative-stain EM studies of HSV-2 gB-B1**.
Representative negative-stain EM 2D class averages for gB-B1 in complex with Fabs 2c, HSV010-13, and BMPC-23.

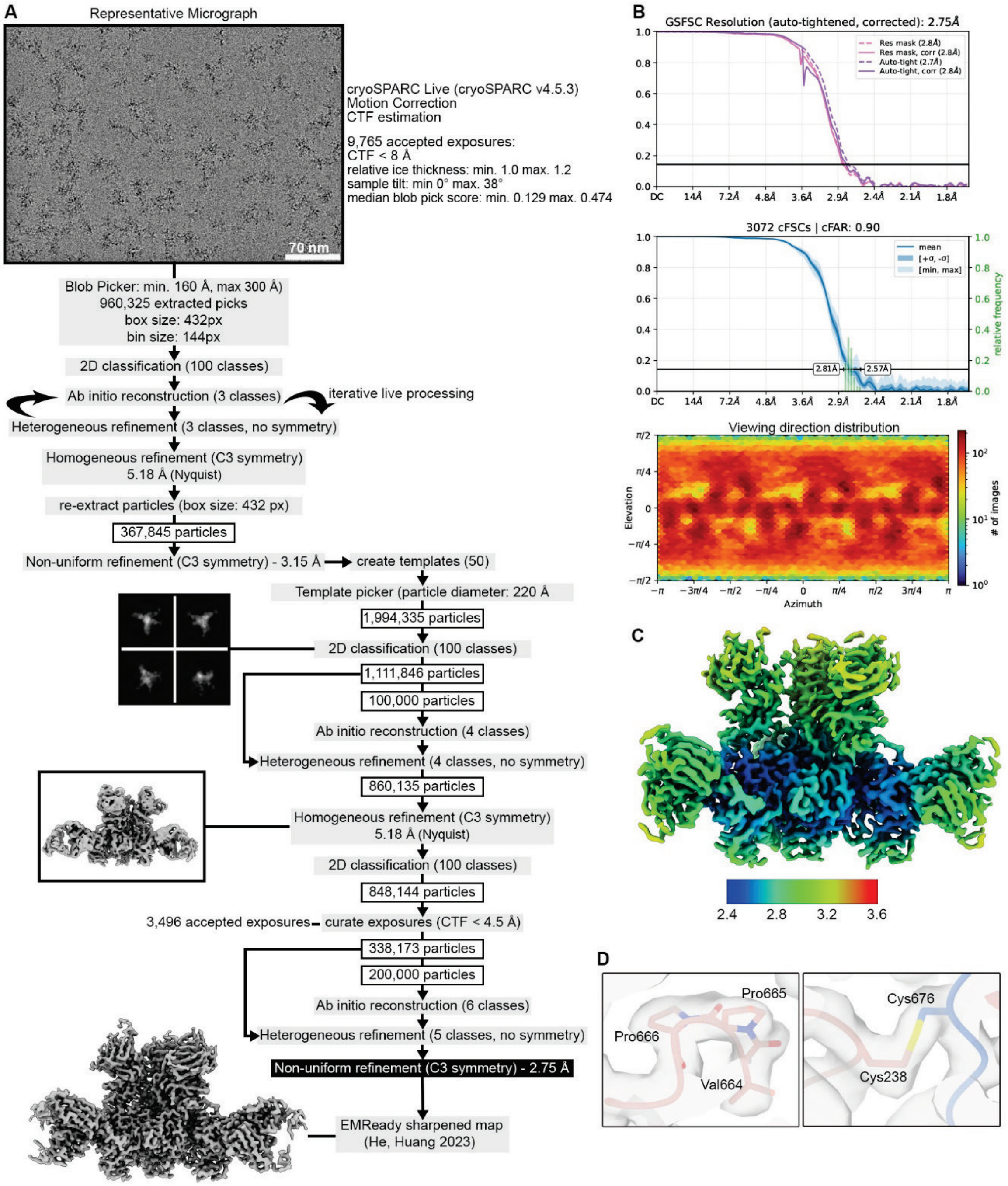


**Supplementary Figure S2: Cryo-EM data processing and evaluation**

**A)** Cryo-EM data processing workflow for the HSV-2 gB-G3 dataset. **B)** GSFSC, cFSC, and viewing-direction distribution plots generated in cryoSPARC. **C)** Unsharpened map produced by non-uniform refinement with C3 symmetry colored by local resolution (Å) using an FSC threshold of 0.143. **D)** EMReady processed map shown as a transparent surface with the HSV-2 gB-G3 structure shown as a ribbon representation with select sidechains shown as sticks. Oxygen atoms are colored red, nitrogen are blue, and sulfur are yellow.

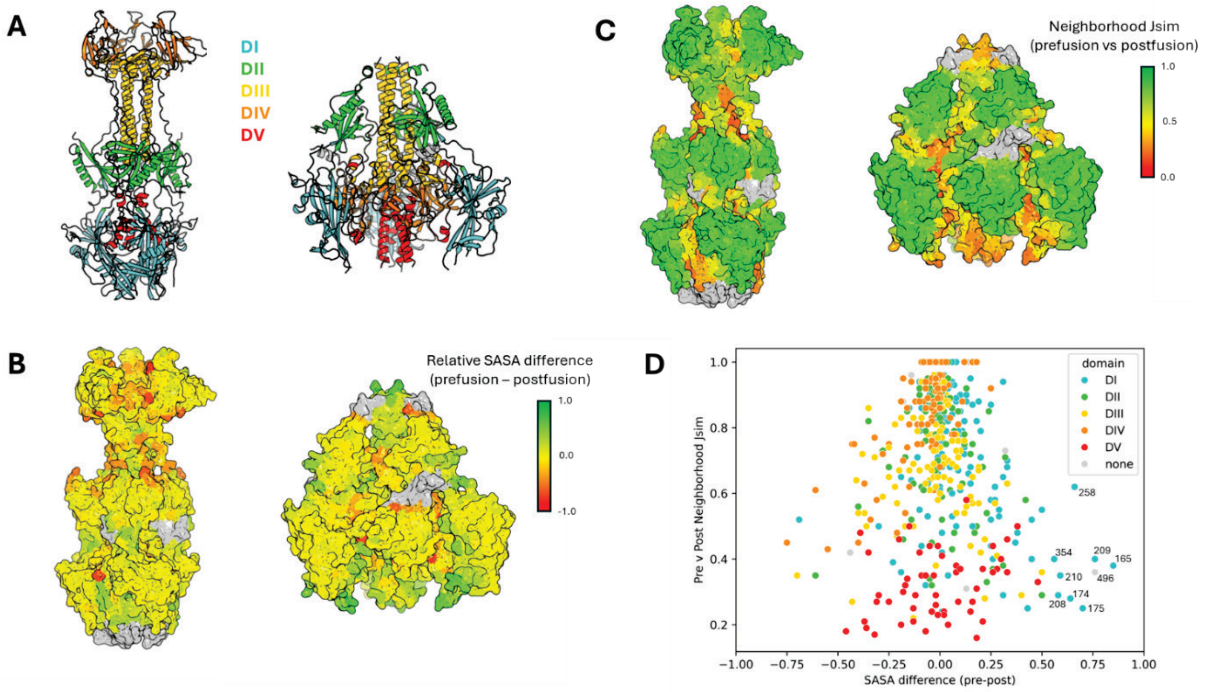


**Supplementary Figure S3: Comparison of Conformational Changes between Prefusion and Postfusion HSV2 gB**.

**A)** Structures of postfusion (PDB 8HR1) and prefusion HSV2 gB (G3, this study) colored by domain. **B)** Relative solvent accessible surface area (SASA) values (0 to 1) were calculated for each residue in both the prefusion and postfusion conformations. Residues in each conformation colored by relative SASA difference (prefusion – postfusion). Green residues are more accessible in the prefusion conformation, and red residues are more accessible in the postfusion conformation. Residues that are not visible in the other conformation are colored gray. **C)** For each residue in each conformation, a neighborhood was defined as all residues within 10 Å of the target residue. Jaccard similarity scores were calculated for neighborhood lists between prefusion and postfusion. Green residues have similar neighborhoods in both conformations and red residues have different neighborhoods in each conformation. **D)** Scatter plot of the per-residue relative SASA difference (x-axis) vs neighborhood Jaccard similarity scores (y-axis) with residues colored by domain. Residues in the bottom right quadrant represent potential prefusion-specific epitopes: those that are both more accessible in the prefusion conformation and have a different neighborhood compared to postfusion.

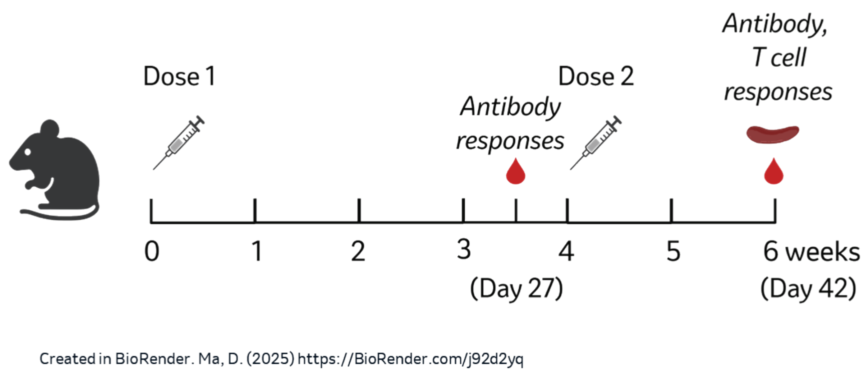


**Supplementary Figure S4: Immunization scheme**
Mice are immunized at week 0 and week 4. Serum samples are collected at day 27 (~4 weeks post-dose 1) and 42 (2 weeks post-dose 2) to characterize antibody responses, and spleens are harvested at day 42 (2 weeks post-dose 2) to characterize T cell responses.

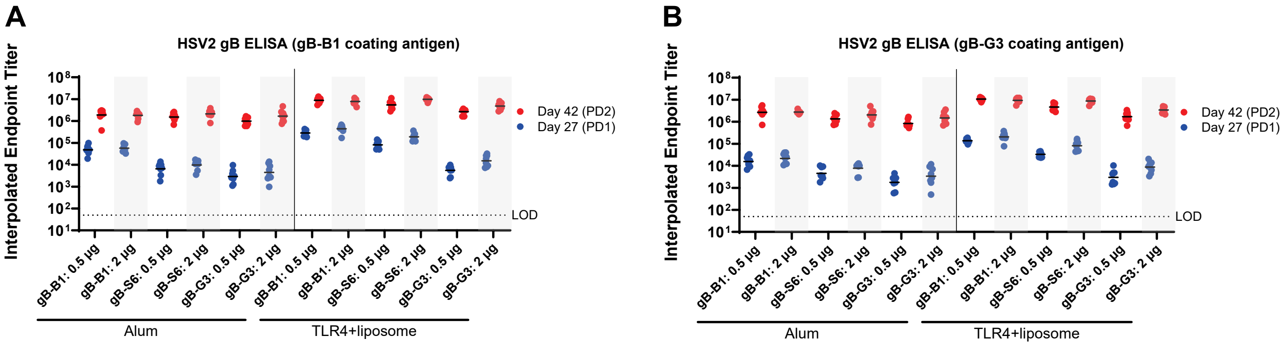


**Supplementary Figure S5. Comparison of coating protein antigen for detection of HSV-2 gB antibody titers by ELISA: Protein immunizations**.

Antibody responses are characterized by ELISA using **A)** postfusion gB-B1 protein or **B)** prefusion gB-G3 as coating antigens to quantify HSV-2 gB binding titers. Symbols represent individual animals (n=8) and bars represent geometric mean.

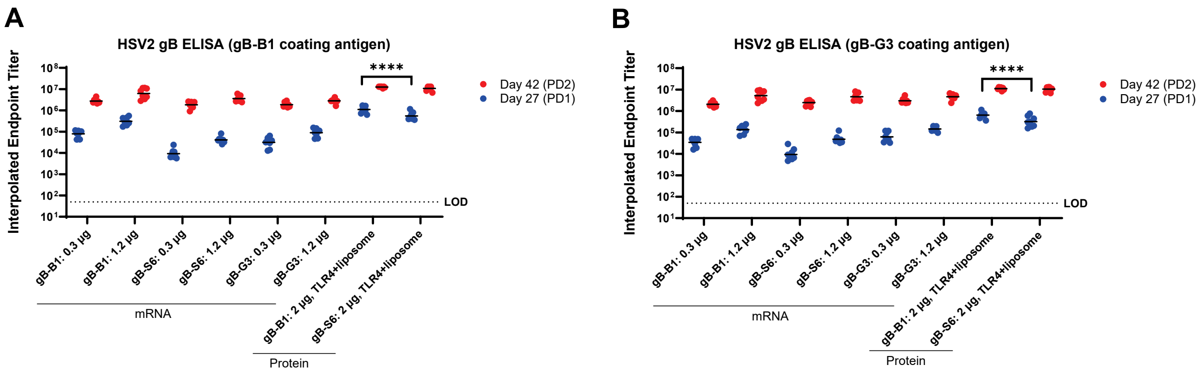


**Supplementary Figure S6. Comparison of coating protein antigen for detection of HSV-2 gB antibody titers by ELISA: mRNA and protein immunizations.**
Antibody responses are characterized by ELISA using **A)** postfusion B1 protein or **B)** prefusion G3 as coating antigens to quantify HSV2 gB binding titers. Symbols represent individual animals (n=7 or 8) and bars represent geometric mean.

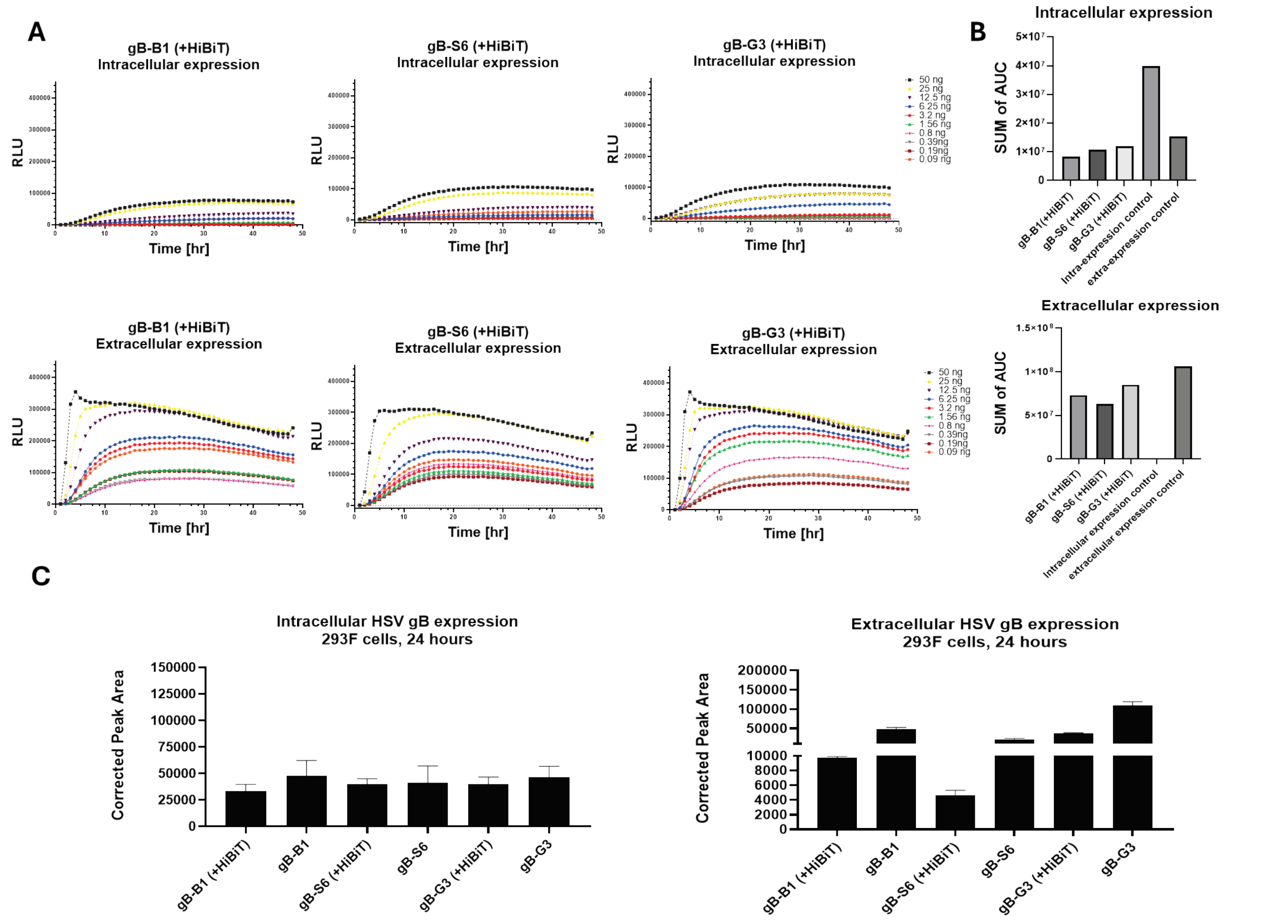


**Supplementary Figure S7.** ***In vitro*** **protein expression from linear mRNA constructs**.

**A)** Expression profiling over 48 hrs of HSV-2 gB variants gB-B1, -S6, and -G3 measured intracellularly (upper panel) and extracellularly (lower panel). **B)** Comparison of HSV-2 gB-B1, -S6, and -G3 expression levels summarized by the overall area under the curve (AUC). Intracelluar expression control: Intracellularly expressed control construct used as a reference for intracellular expression. Extracellular expression control: Secreted control construct used as a reference for extracellular expression. **C**) Western blot quantification of (left) intracellular or (right) extracellular HSV gB expression at 24 hrs post-transfection of 293F cells with linear mRNA constructs with and without HiBiT tags. Bar graphs show the mean corrected peak area and standard deviation from two independent experiments.

**Supplementary Table 1. Data collection and refinement statistics**

| | HSV-2 prefusion gB-G3 |
|---|---|
| **Data collection** | |
| Microscope | FEI Titan Krios |
| Voltage (kV) | 300 |
| Detector | Gatan K3 |
| Magnification (nominal) | 105,000 |
| Pixel size (Å/pix) | 0.8332 |
| Exposure rate ($e^-$/pix/sec) | 11 |
| Exposure dose ($e^-$/Å$^2$) | 70 |
| Defocus range (μm) | 1.0–3.0 |
| Tilt angle (°) | 0 and 30 |
| Micrographs collected | 11,000 |
| Micrographs used | 9,765 |
| Automation software | SerialEM |
| **Data processing** | |
| Particles | 227,517 |
| Symmetry | C3 |
| Map sharpening B-factor | 93.0 |
| Resolution (Å) at FSC (half-maps) | |
| Unmasked: 0.5 | 3.5 |
| Masked: 0.5 | 3.1 |
| Unmasked: 0.143 | 3.2 |
| Masked: 0.143 | 2.75 |
| EMDB ID | XXXXX |
| **Model Refinement** | |
| Composition | |
| Amino Acids | 3,099 |
| Ligands (type) | - |
| Bonds (RMSD) | |
| Length (Å) | 0.003 |
| Angles (°) | 0.63 |
| Ramachandran plot | |
| Outliers (%) | 0.0 |
| Allowed (%) | 4.6 |
| Favored (%) | 95.4 |
| Rotamer outliers (%) | 0.0 |
| C-β outliers (%) | 0.0 |
| CaBLAM outliers (%) | 2.1 |
| ADP (B-factors) | |
| Amino Acids (mean) | 55.4 |
| Ligands (mean) | - |
| MolProbity score | 1.68 |
| Clash score | 6.49 |
| PDB ID | XXXX |

### Supplementary Table 2. GMTs, p-values and fold change comparisons for antibody responses: Protein immunizations

| Adjuvant | Vaccine antigen | Protein dose (µg) | Geometric mean titers (GMT) | Geometric Standard deviation | n | p-values: gB-B1 vs Pre-fusion GMTs (Protein dose and adjuvant matched) | p-values: S6 vs G3 GMTs (Protein dose and adjuvant matched) | Fold change: TLR4+liposome/ Alum (Protein dose and adjuvant matched) | Fold change: gB-B1/Pre-fusion (Protein dose and adjuvant matched) | Fold change: S6/G3 (Protein dose and adjuvant matched) | Geometric mean | Geometric Standard deviation | n | p-values: gB-B1 vs Pre-fusion GMTs (Protein dose and adjuvant matched) | p-values: S6 vs G3 GMTs (Protein dose and adjuvant matched) | Fold change: TLR4+liposome/ Alum (Protein dose and adjuvant matched) | Fold change: gB-B1/Pre-fusion (Protein dose and adjuvant matched) | Fold change: S6/G3 (Protein dose and adjuvant matched) |
|---|---|---|---|---|---|---|---|---|---|---|---|---|---|---|---|---|---|---|
| Alum | gB-B1 | 0.5 | 4.80E+04 | 1.6 | 8 | - | - | - | - | - | 1.88E+06 | 2.0 | 8 | - | - | - | - | - |
| | gB-B1 | 2 | 5.73E+04 | 1.5 | 8 | - | - | - | - | - | 1.81E+06 | 1.4 | 8 | - | - | - | - | - |
| | gB-S6 | 0.5 | 6.55E+03 | 2.0 | 8 | ns | - | - | 7.3 | - | 1.53E+06 | 1.5 | 8 | ns | - | - | 1.2 | - |
| | gB-S6 | 2 | 9.77E+03 | 1.7 | 8 | ns | - | - | 5.9 | - | 2.14E+06 | 1.6 | 8 | ns | - | - | 0.8 | - |
| | gB-G3 | 0.5 | 2.92E+03 | 2.0 | 8 | ns | ns | - | 16.4 | 2.2 | 9.93E+05 | 1.5 | 8 | ns | ns | - | 1.9 | 1.5 |
| | gB-G3 | 2 | 4.50E+03 | 2.5 | 8 | ns | ns | - | 12.7 | 2.2 | 1.67E+06 | 1.8 | 8 | ns | ns | - | 1.1 | 1.3 |
| TLR4+ liposome | gB-B1 | 0.5 | 2.86E+05 | 1.4 | 8 | - | - | 5.9 | - | - | 8.83E+06 | 1.3 | 8 | - | - | 4.7 | - | - |
| | gB-B1 | 2 | 4.42E+05 | 1.5 | 8 | - | - | 7.7 | - | - | 7.84E+06 | 1.3 | 8 | - | - | 4.3 | - | - |
| | gB-S6 | 0.5 | 8.20E+04 | 1.5 | 8 | **** | - | 12.5 | 3.5 | - | 5.48E+06 | 1.5 | 8 | ** | - | 3.6 | 1.6 | - |
| | gB-S6 | 2 | 1.90E+05 | 1.5 | 8 | **** | - | 19.4 | 2.3 | - | 9.82E+06 | 1.2 | 8 | ns | - | 4.6 | 0.8 | - |
| | gB-G3 | 0.5 | 5.57E+03 | 1.7 | 8 | **** | ns | 1.9 | 51.3 | 14.7 | 2.69E+06 | 1.4 | 8 | **** | ** | 2.7 | 3.3 | 2.0 |
| | gB-G3 | 2 | 1.51E+04 | 1.8 | 8 | **** | **** | 3.4 | 29.2 | 12.5 | 4.81E+06 | 1.4 | 8 | ** | **** | 2.9 | 1.6 | 2.0 |

| Serum neutralization titers: Protein Immunizations | | | Day 27 | | | | | | | | Day 42 | | | | | | | |
|---|---|---|---|---|---|---|---|---|---|---|---|---|---|---|---|---|---|---|
| Adjuvant | Vaccine antigen | Protein dose (µg) | Geometric mean titers (GMT) | Geometric Standard deviation | n | p-values: gB-B1 vs Pre-fusion GMTs (Protein dose and adjuvant matched) | p-values: S6 vs G3 GMTs (Protein dose and adjuvant matched) | Fold change: TLR4+liposome/ Alum (Protein dose and adjuvant matched) | Fold change: gB-B1/Pre-fusion (Protein dose and adjuvant matched) | Fold change: S6/G3 (Protein dose and adjuvant matched) | Geometric mean | Geometric Standard deviation | n | p-values: gB-B1 vs Pre-fusion GMTs (Protein dose and adjuvant matched) | p-values: S6 vs G3 GMTs (Protein dose and adjuvant matched) | Fold change: TLR4+liposome/ Alum (Protein dose and adjuvant matched) | Fold change: gB-B1/Pre-fusion (Protein dose and adjuvant matched) | Fold change: S6/G3 (Protein dose and adjuvant matched) |
| Alum | gB-B1 | 0.5 | 2.00E+01 | 1.0 | 8 | - | - | - | - | - | 1.55E+02 | 2.3 | 8 | - | - | - | - | - |
| | gB-B1 | 2 | 2.19E+01 | 1.2 | 8 | - | - | - | - | - | 3.10E+02 | 4.0 | 8 | - | - | - | - | - |
| | gB-S6 | 0.5 | 2.00E+01 | 1.0 | 8 | ns | - | - | 1.0 | - | 1.86E+02 | 2.2 | 8 | ns | - | - | 0.8 | - |
| | gB-S6 | 2 | 2.00E+01 | 1.0 | 8 | ns | - | - | 1.1 | - | 3.99E+02 | 2.9 | 8 | ns | - | - | 0.8 | - |
| | gB-G3 | 0.5 | 2.00E+01 | 1.0 | 8 | ns | ns | - | 1.0 | 1.0 | 4.00E+01 | 2.8 | 8 | ns | ns | - | 3.9 | 4.7 |
| | gB-G3 | 2 | 2.00E+01 | 1.0 | 8 | ns | ns | - | 1.1 | 1.0 | 6.35E+01 | 2.0 | 8 | ns | ns | - | 4.9 | 6.3 |
| TLR4+ liposome | gB-B1 | 0.5 | 3.55E+02 | 1.7 | 8 | - | - | 17.7 | - | - | 4.17E+04 | 1.5 | 8 | - | - | 269.0 | - | - |
| | gB-B1 | 2 | 4.81E+02 | 2.1 | 8 | - | - | 21.9 | - | - | 3.98E+04 | 1.7 | 8 | - | - | 128.6 | - | - |
| | gB-S6 | 0.5 | 7.55E+01 | 2.0 | 8 | **** | - | 3.8 | 4.7 | - | 2.16E+04 | 1.9 | 8 | ** | - | 116.0 | 1.9 | - |
| | gB-S6 | 2 | 2.90E+02 | 2.0 | 8 | * | - | 14.5 | 1.7 | - | 3.39E+04 | 1.6 | 8 | ns | - | 84.7 | 1.2 | - |
| | gB-G3 | 0.5 | 2.00E+01 | 1.0 | 8 | **** | ns | 1.0 | 17.7 | 3.8 | 1.01E+03 | 5.8 | 8 | **** | *** | 25.2 | 41.4 | 21.4 |
| | gB-G3 | 2 | 2.00E+01 | 1.0 | 8 | **** | **** | 1.0 | 24.1 | 14.5 | 4.10E+03 | 2.4 | 8 | **** | **** | 64.6 | 9.7 | 8.2 |

Geometric mean titers, statistics and fold change comparisons for antibody responses in protein immunization study. Where available, asterisks represent statistically significant differences for comparisons of geometric mean titers (GMTs) between two groups where "-" = not determined; ns = not significant; *p<0.05; **p<0.01; ***p<0.001 and ***

### Supplementary Table 3. GMTs, p-values and fold change comparisons for antibody responses: mRNA and protein immunizations

| ELISA titers: mRNA and Protein Immunizations | | | Day 27 | | | | | | | | Day 42 | | | | | | | |
|---|---|---|---|---|---|---|---|---|---|---|---|---|---|---|---|---|---|---|
| Vaccine modality | Vaccine antigen | Vaccine dose (mRNA or Protein, μg) | Geometric mean titers (GMT) | Geometric Standard deviation | n | p-values: gB-B1 vs Pre-fusion GMTs (Antigen dose matched) | p-values: S6 vs G3 mRNA GMTs (Antigen dose matched) | Fold change: mRNA gB-B1/Pre-fusion (Dose matched) | Fold change: mRNA S6/G3 (Dose matched) | Fold change: Protein B1/S6 | Geometric mean | Geometric Standard deviation | n | p-values: gB-B1 vs Pre-fusion GMTs (Antigen dose matched) | p-values: S6 vs G3 mRNA GMTs (Antigen dose matched) | Fold change: mRNA gB-B1/Pre-fusion (Dose matched) | Fold change: mRNA S6/G3 (Dose matched) | Fold change: Protein B1/S6 |
| mRNA | gB-B1 | 0.3 | 7.93E+04 | 1.5 | 8 | - | - | - | - | - | 2.74E+06 | 1.3 | 8 | - | - | - | - | - |
| | gB-B1 | 1.2 | 3.09E+05 | 1.6 | 7 | - | - | - | - | - | 6.26E+06 | 1.8 | 8 | - | - | - | - | - |
| | gB-S6 | 0.3 | 9.23E+03 | 1.6 | 8 | ns | - | 8.6 | - | - | 1.84E+06 | 1.4 | 8 | ns | - | 1.5 | - | - |
| | gB-S6 | 1.2 | 4.11E+04 | 1.4 | 8 | ns | - | 7.5 | - | - | 3.58E+06 | 1.4 | 8 | ** | - | 1.7 | - | - |
| | gB-G3 | 0.3 | 3.13E+04 | 1.8 | 8 | ns | ns | 2.5 | 0.3 | - | 1.86E+06 | 1.3 | 8 | ns | ns | 1.5 | 1.0 | - |
| | gB-G3 | 1.2 | 8.94E+04 | 1.6 | 8 | ns | ns | 3.5 | 0.5 | - | 2.81E+06 | 1.3 | 8 | *** | ns | 2.2 | 1.3 | - |
| Protein | gB-B1 | 2 | 1.11E+06 | 1.4 | 8 | - | - | - | - | - | 1.26E+07 | 1.1 | 8 | - | - | - | - | - |
| | gB-S6 | 2 | 5.46E+05 | 1.5 | 8 | **** | - | - | - | 2.0 | 1.08E+07 | 1.3 | 8 | ns | - | - | - | 1.2 |

| Serum neutralization titers: mRNA and Protein Immunizations | | | Day 27 | | | | | | | | Day 42 | | | | | | | |
|---|---|---|---|---|---|---|---|---|---|---|---|---|---|---|---|---|---|---|
| Vaccine modality | Vaccine antigen | Vaccine dose (mRNA or Protein, μg) | Geometric mean titers (GMT) | Geometric Standard deviation | n | p-values: gB-B1 vs Pre-fusion GMTs (Antigen dose matched) | p-values: S6 vs G3 mRNA GMTs (Antigen dose matched) | Fold change: mRNA gB-B1/Pre-fusion (Dose matched) | Fold change: mRNA S6/G3 (Dose matched) | Fold change: Protein B1/S6 | Geometric mean | Geometric Standard deviation | n | p-values: gB-B1 vs Pre-fusion GMTs (Antigen dose matched) | p-values: S6 vs G3 mRNA GMTs (Antigen dose matched) | Fold change: mRNA gB-B1/Pre-fusion (Dose matched) | Fold change: mRNA S6/G3 (Dose matched) | Fold change: Protein B1/S6 |
| mRNA | gB-B1 | 0.3 | 4.53E+01 | 1.4 | 8 | - | - | - | - | - | 2.55E+03 | 1.8 | 8 | - | - | - | - | - |
| | gB-B1 | 1.2 | 2.17E+02 | 1.8 | 7 | - | - | - | - | - | 5.79E+03 | 1.5 | 8 | - | - | - | - | - |
| | gB-S6 | 0.3 | 2.00E+01 | 1.0 | 8 | ns | - | 2.3 | - | - | 6.39E+02 | 1.9 | 8 | ns | - | 4.0 | - | - |
| | gB-S6 | 1.2 | 2.51E+01 | 1.6 | 8 | ns | - | 8.7 | - | - | 1.81E+03 | 7.0 | 8 | ns | - | 3.2 | - | - |
| | gB-G3 | 0.3 | 2.36E+01 | 1.4 | 8 | ns | ns | 1.9 | 0.8 | - | 1.86E+03 | 1.7 | 8 | ns | ns | 1.4 | 0.3 | - |
| | gB-G3 | 1.2 | 5.81E+01 | 1.4 | 8 | ns | ns | 3.7 | 0.4 | - | 3.50E+03 | 2.6 | 8 | ns | ns | 1.7 | 0.5 | - |
| Protein | gB-B1 | 2 | 1.34E+03 | 2.2 | 8 | - | - | - | - | - | 2.49E+04 | 2.0 | 8 | - | - | - | - | - |
| | gB-S6 | 2 | 4.97E+02 | 2.4 | 8 | ** | - | - | - | 2.7 | 2.48E+04 | 2.2 | 8 | ns | - | - | - | 1.0 |

Geometric mean titers, statistics and fold change comparisons for antibody responses in mRNA and protein immunization study. Where available, asterisks represent statistically significant differences for comparisons of geometric mean titers (GMTs) between two groups where "-" = not determined; ns = not significant; **p<0.01; ***p<0.001 and ****p<0.0001.
